# Restoration of the Functional *nif* Gene Cluster by Seven Excision and Two Inversion Events during Heterocyst Development in the Nitrogen-Fixing Cyanobacterium *Calothrix* sp. NIES-4101

**DOI:** 10.1101/2023.12.11.571196

**Authors:** Kazuma Uesaka, Mari Banba, Sotaro Chiba, Yuichi Fujita

**Author notes:** Corresponding Author: Y. Fujita Graduate School of Bioagricultural Sciences, Nagoya University Furo-cho, **Chikusa-ku**, Nagoya **464-8601 **. equally contributed (Kazuma Uesaka, Mari Banba).

## Abstract

In the genome of the heterocystous cyanobacterium *Calothrix* sp. NIES-4101 (NIES-4101), the four genes essential for nitrogen fixation (*nifB*, *nifH*, *nifD*, and *nifK*) are highly fragmented into 13 parts in a 350-kb chromosomal region, and four of these parts are encoded in the reverse strand. Such a complex fragmentation feature makes it difficult to restore the intact *nifBHDK* genes by the excision mechanism found in the *nifD* gene of the *Anabaena* sp. PCC 7120 heterocyst. To examine the nitrogen-fixing ability of NIES-4101, we confirmed that NIES-4101 grew well on combined nitrogen-free medium and showed high nitrogenase activity, which strongly suggested that the complete *nifBHDK* genes are restored by a complex recombination process in heterocysts. Next, we resequenced the genome prepared from cells grown under nitrogen-fixing conditions. Two contigs covering the complete *nifHDK* and *nifB* genes were found by *de novo* assembly of the sequencing reads. In addition, DNA fragments covering the *nifBHDK* operon were successfully amplified by PCR. We propose that the process of *nifBHDK* restoration occurs as follows. First, the *nifD-nifK* genes are restored by four excision events. Then the complete *nifH* and *nifB* genes are restored by two excision events followed by two successive inversion events between the inverted repeat sequences and one excision event, forming the functional *nif* gene cluster, *nifB-fdxN-nifS-nifU-nifH-nifD-nifK*. All genes coding recombinases responsible for these nine recombination events are located close to the terminal repeat sequences. The restoration of the *nifBHDK* genes in NIES-4101 is the most complex genome reorganization reported in heterocystous cyanobacteria.

## Introduction

Nitrogen fixation is the process of converting dinitrogen (N_2_) into ammonia. Nitrogenase, which catalyzes the nitrogen fixation reaction, is extremely sensitive to oxygen (Yates and Planqué 1975). Thus, nitrogen-fixing bacteria growing in aerobic environments require a variety of molecular mechanisms to protect nitrogenase from oxygen (Fay 1992, Fujita and Uesaka 2022). Cyanobacteria perform the oxygenic photosynthesis in the same way as plants, and approximately half of cyanobacterial species have the ability of nitrogen fixation (Stal and Zehr 2008). Since cyanobacteria evolve oxygen through photosynthesis, nitrogen-fixing cyanobacteria must solve the “oxygen paradox” to accommodate nitrogenase in cells carrying out oxygenic photosynthesis. The best-known of these protection mechanisms is the spatial separation of nitrogenase from photosynthesis by differentiation of nitrogen-fixing cells, heterocysts, which are differentiated from vegetative cells (Kumar et al. 2010, Maldener et al. 2014, Zeng and Zhang 2022). Heterocysts suppress the activity of photosystem II, which evolves oxygen (Sugiura et al. 2012), and produce a thick cell wall that prevents the influx of oxygen from outside the cell (Awai and Wolk 2007). The small amount of oxygen that penetrates the cell wall is removed by the high respiratory activity of the cell membrane (Valladares et al. 2003). These oxygen protection mechanisms create hypoxic environments in the heterocysts so that nitrogenase can operate, allowing the cyanobacteria to grow by nitrogen fixation even in aerobic environments.

Most studies into heterocyst differentiation have been carried out using *Anabaena* sp. PCC 7120 (*A.* 7120) (Golden and Yoon 2003, Herrero and Flores 2019). In nitrogen-deficient conditions, *A.* 7120 vegetative cells differentiate into heterocysts at a rate of approximately 1 in every 8 to 15 vegetative cells at semi-regular intervals (Yoon and Golden 2001). The process of heterocyst differentiation is a complex cascade involving nearly 600–1,000 genes (Lynn et al. 1986, Adams and Duggan 1999, Flores et al. 2019), which is mainly regulated by the transcriptional regulatory protein HetR (Buikema and Haselkorn 1991, Black et al. 1993, Kim et al. 2013), and takes around 20–24 h to be complete. One of the most important findings in early studies of heterocyst differentiation in *A.* 7120 is that the excision of the 11-kb insertion sequence that disrupts the *nifD* coding frame by a site-specific recombinase (XisA) during heterocyst differentiation to restore the intact *nifD* gene (Golden et al. 1985, Lammers et al. 1986, Golden and Wiest 1988). Similar excision events of insertion sequences have been reported for *fdxN*, encoding ferredoxin (Carrasco et al. 1994), and *hupL*, encoding the large subunit of uptake-hydrogenase (Carrasco et al. 1995, Carrasco et al. 2005). The presence of insertion sequences that interrupt the coding frames of *nif* genes in the genomes of 38 heterocyst-forming cyanobacterial species has been systematically investigated (Hilton et al. 2016). A total of 101 interruption elements were defined in a wide variety of genes, including *nifH*, *nifB*, *nifE* and *nifJ* as well as *nifD*, *fdxN*, and *hupL*.

While heterocyst-forming cyanobacteria are monophyletic (Giovannoni et al. 1988, Howard-Azzeh et al. 2014), they form a diverse cyanobacterial group with different morphological characteristics (heterocyst formation pattern, hormogonia formation, tapering, branching, etc.) and a variety of growth environments (freshwater, land, ocean, hot springs, etc.). The genus *Calothrix* develop heterocysts at the terminal cells of trichomes and have a morphological characteristic in which the trichomes taper from the end with the heterocyst toward the tip. It has been reported that the presence of fragmented *nif* genes is widely distributed in heterocyst-forming cyanobacteria, including *Calothrix* (Hilton et al. 2016). However, considering that *Calothrix* has a significantly different heterocyst formation pattern from *A.* 7120 and the *Calothrix* genus is evolutionarily distant from the *Anabaena* genus including *A.* 7120 in phylogenetic trees (Howard-Azzeh et al. 2014), we expect that *Calothrix* exhibits different characteristics from *A.* 7120 in the reconstruction of fragmented *nif* genes during heterocyst differentiation. The features of the fragmentation and the reconstruction of the *nif* genes may provide a new perspective on the evolution of heterocyst-forming cyanobacteria.

DNA rearrangements during cell differentiation and for the transcriptional regulation of genes are rare in prokaryotes, except for heterocystous cyanobacteria. One example of genome rearrangement during cell differentiation in prokaryotes is the *sigK* gene encoding the RNA polymerase sigma factor, which is reconstituted by the site-directed recombinase CisA by the excision of the 42-kb insertion sequence during spore formation in *Bacillus subtilis* (Stragier et al. 1989, Kunkel et al. 1990). The insertion sequence is flanked by direct repeat sequences that are recognized and excised by CisA. It has also been reported that gene expression is regulated by the inversion of the insertion sequence between short inverted repeats through the action of a recombinase in *Salmonella typhimurium* (Zieg et al. 1977, Zieg and Simon 1980, Johnson and Simon 1985).

In this study, we focused on the heterocyst-forming filamentous cyanobacterium *Calothrix* sp. NIES-4101 (NIES-4101), whose complete genome sequence was recently determined by Hirose *et al* (Hirose et al. 2021). When we examined the *nif* gene cluster in the genome of NIES-4101, we noticed that the four genes (*nifB*, *nifH*, *nifD*, and *nifK*), which are essential for nitrogen fixation, were highly fragmented. Surprisingly, the coding regions of the *nifBHDK* genes are fragmented into 13 parts in a 350-kb chromosomal region: four parts for *nifH* (*nifH*-I, II, III, and IV) and *nifD* (*nifD*-I, II, III, and IV), two for *nifK* (*nifK*-I and II), and three for *nifB* (*nifB*-I, II, and III). Furthermore, four of these parts (*nifH*-III, *nifB*-III, *nifH*-I, and *nifH*-II) are encoded in the reverse strand. In addition, the arrangement of the four parts of *nifH* and the three parts of *nifB* is highly mixed along the chromosomal region: *nifH*-IV, *nifB*-II, *nifB*-I, *nifH*-III, *nifH*-I, and *nifH*-II. Therefore, a simple repeat of multiple excision events as found in *A.* 7120 will never give rise to the complete coding frames of the *nifBHDK* genes in heterocysts of NIES-4101.

To confirm the nitrogen fixation capability of NIES-4101, we first examined its diazotrophic growth and nitrogenase activity. We then resequenced genomic DNA extracted from NIES-4101 cells grown under nitrogen fixation conditions. Two contigs covering the full-length *nifHDK* and *nifB* genes were detected at a low rate. The presence of the full-length *nifBHDK* genes was also confirmed by PCR. The results indicated that the 13 parts of the four *nifHDKB* genes are precisely rearranged to create a well-conserved *nif* gene cluster (*nifB-fdxN-nifS-nifU-nifH-nifD-nifK-nifE-nifN-nifX*) in the heterocyst cells. We propose that the restoration process involves six excision events followed by two sequential inversion events between the short inverted repeats, and finally one excision event. The restoration process consisting of seven excision and two inversion events is the most complex gene reorganization known to occur during heterocyst differentiation.

## Results

### Highly fragmented nif genes in the Calothrix sp. NIES-4101 genome

Six *nif* genes (*nifH*, *nifD*, *nifK*, *nifE*, *nifN*, *nifB*) are proposed to comprise the minimal unit essential for nitrogen fixation. The current GenBank annotation of the NIES-4101 complete genome assembly (GCA_004296455.1) has *nifE* (NIES-4101_63770) and *nifN* (NIES-4101_63760) located next to each other as complete coding frames. However, the other four genes, *nifH*, *nifD*, *nifK*, and *nifB*, are highly fragmented into 4, 4, 2, and 3 parts (total 13), respectively, in the about 350-kb region located next to the *nifE* and *nifN* genes (Fig. 1A). To rule out the possibility of misassembly of the genome, the sequencing reads (DRR315835) were downloaded from the Sequence Read Archive (SRA) and analyzed carefully. No assembly errors were detected in the *nif* locus, including other regions. We concluded that the fragmentation of the *nifBHDK* genes was not an artifact of the genome assembly.

**Figure 1.**
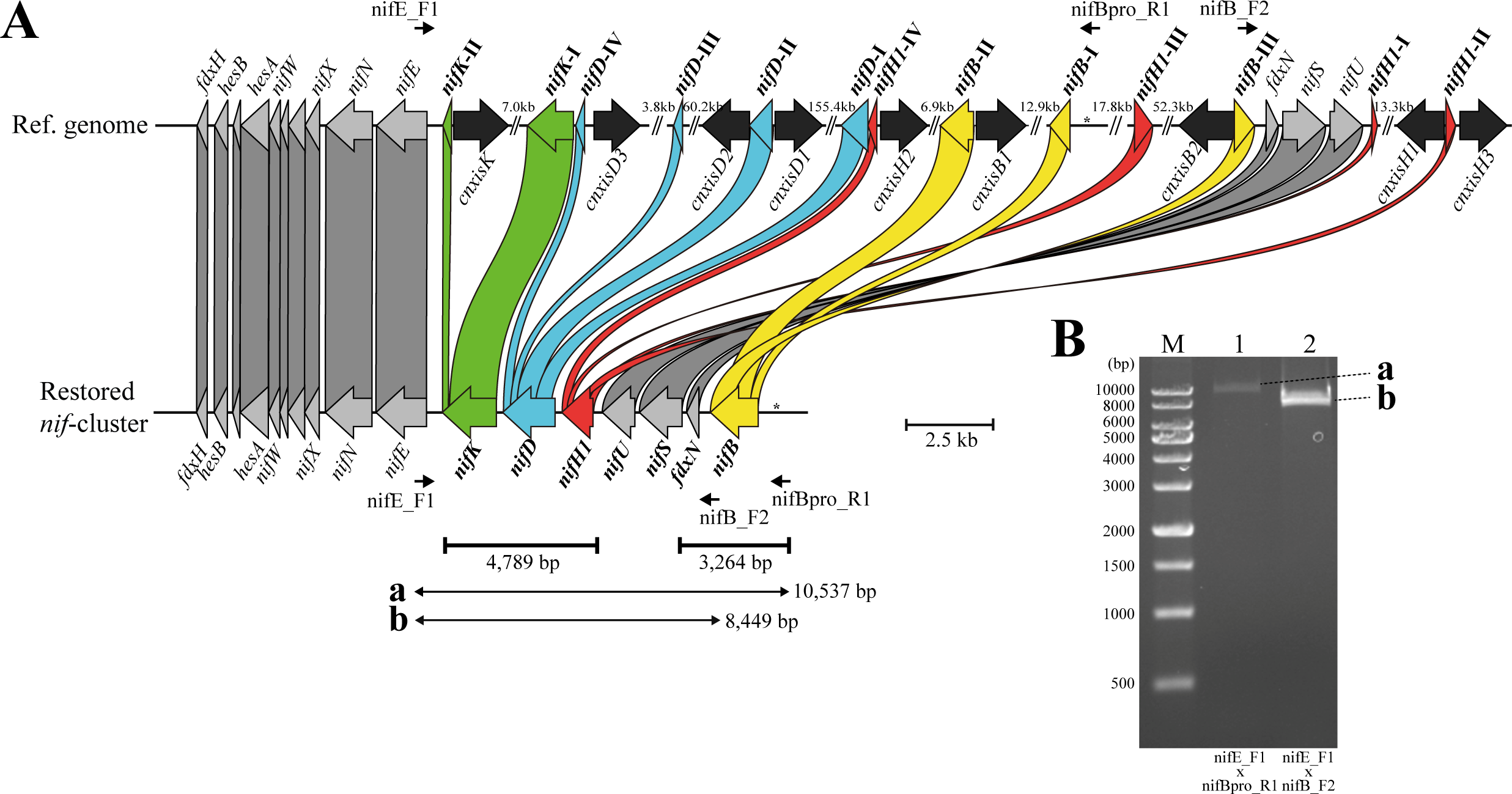
Fragmentation feature of *nifBH1DK* genes in genomes of vegetative cells and the restored *nif* gene cluster in the heterocyst genome of NIES-4101. (**A**) The 350-kb chromosome region of vegetative cells (top) is restored as the 17-kb *nif* gene cluster (bottom) by a series of excision and inversion events. Thirteen fragmented parts of the four genes *nifB*, *nifH1*, *nifD*, and *nifK* are scattered across the 350-kb region. The colored box represents the 13 fragmented parts of the *nifBH1DK* genes in the genome of vegetative cells. Black colored arrow-boxes represent the nine genes encoding site-specific recombinases. Contigs assembled from rearrangement junction sequencing reads are shown as thick horizontal bars with their size. Horizontal bars (**a** and **b**) with arrows at both ends with the fragment size indicate the regions corresponding to the PCR fragments. The long insertion sequences are indicated by “//”. The primers (nifE_F1, nifBpro_R1, and nifB_F2) used for PCR are shown by short arrows (Supplementary Table S5). Asterisk represents the *cis* element in the *nifB* promoter. This figure was drawn using Clinker (Gilchrist and Chooi 2021). (**B**) PCR results detecting two DNA fragments covering the restored *nifBH1DK* region. The signals, **a** (lane 1) and **b** (lane 2), are corresponding to the horizontal bars **a** and **b**, respectively in panel **A**. Lane M is a molecular size marker (XL DNA Ladder 1K, Apro Science, Tokushima).

### Fragmentation feature of nifHDKB

For *nifH*, two *nifH* genes are found in the NIES-4101 genome. One *nifH* gene is a full-length coding open reading frame (ORF) and the other *nifH* gene is four-part fragmented in the 350-kb region (Table 1, Fig. 1A). The full-length *nifH* coding ORF is present as a monocistronic gene approximately 2.76-Mb away from the *nif* gene cluster. Given that a monocistronic *nifH* has been reported as a gene with unknown function in *A.* 7120 (Kumar et al. 2019), the four-part fragmented *nifH* gene that is located within the region of the *nif* gene cluster was designated *nifH1* as an essential gene for nitrogen fixation, and the other full-length *nifH* gene was designated *nifH2* (Kumar et al. 2019). The features of the *nifH1* fragmentation are as follows (Fig. 1A): four fragments are scattered in a 115-kb region, with a maximum distance of 58 kb between fragments. Based on the homology to the *A.*7120 NifH protein (AAA22008.1), the first part (*nifH1*-I for the N-terminal 1–48 amino acid sequence) is located adjacent to the *nifU* gene, the second part (*nifH1*-II for the central 48–139 amino acid sequence) is 13.3-kb downstream of the *nifH1*-I, the third part (*nifH1*-III for the central 135–264 amino acid sequence) is located between the *nifB*-I and *nifB*-III, and the fourth part (*nifH1*-IV for the C-terminal 260–296 amino acid sequence) is located at adjacent to the *nifD*-I part (Table 1). In addition, only *nifH1*-IV is oriented in the same direction as *nifEN* and *nifD*-I, while the other three are oriented in the opposite direction. Due to the inconsistent coding order and direction, the full-length *nifH1* cannot be restored by simple excision of the insertion sequences between *nifH1* fragments.

**Table 1.** *nifHDKENB* coding regions in NIES-4101 vegetative genome assembly.

| Gene | Copy Number | Full length | Fragmented * |
| --- | --- | --- | --- |
| *nifH* | 2 | *nifH2*<br>+:1,620,269–1,621,153 (1..295 aa) | *nifH1*<br>–: 4,384,791–4,384,898 (260..295 aa) (IV)<br>+: 4,426,358–4,426,739 (135..264 aa) (III)<br>+: 4,484,564–4,484,708 (1..48 aa) (I)<br>+: 4,499,311–4,499,589 (48..139 aa) (II) |
| *nifD* | 1 | Not found | –: 4,159,942–4,160,078 (451..494 aa) (IV)<br>–: 4,164,808–4,164,853 (439–453 aa) (III)<br>–: 4,226,413–4,227,024 (235..440 aa) (II)<br>–: 4,383,892–4,384,602 (1..237 aa) (I) |
| *nifK* | 1 | Not found | –: 4,149,612–4,149,866 (426..511 aa) (II)<br>–: 4,158,486–4,159,765 (1..426 aa) (I) |
| *nifE* | 1 | –:4,147,777–4,149,162 (1..463 aa) | Not found |
| *nifN* | 1 | –:4,146,311–4,147,615 (1..435 aa) | Not found |
| *nifB* | 1 | Not found | –: 4,393,274–4,394,161 (26–321 aa) (II)<br>–: 4,408,459–4,408,548 (1..30 aa) (I)<br>+: 4,480,730–4,481,240 (317..485 aa) (III) |

\* +/- indicates the orientation. Exact coding sequence lengths are unknown due to genomic rearrangement events with several base-pair overlaps. Roman numerals in parentheses correspond to the notations in Figures 1 and 4.

The *nifD* gene is fragmented into four parts (*nifD*-I, II, III, and IV) that span over 225 kb, and all fragments are in the same direction. The maximum insertion sequence is 157 kb in length and located between *nifD*-II and *nifD*-I. The *nifK* gene is fragmented into two parts, with one part encoding the N-terminal 1–426 amino acid residues (*nifK*-I) and the other part encoding the C-terminal 426–513 amino acid residues (*nifK*-II). The two fragments are separated by 8.6 kb and are both are in the same direction as the *nifD* parts. The full-length *nifD* and *nifK* genes can be restored by simple excision of the insertion sequences (three for *nifD* and one for *nifK*).

The *nifB* gene is fragmented into three parts in a region spanning over 88 kb (Fig. 1A). The three parts are encoded in the order of a central 26–321 amino acid sequence (*nifB*-II), an N-terminal 1–30 amino acid sequence (*nifB*-I), and a C-terminal 317–485 amino acid sequence (*nifB*-III). Due to the opposite directions of *nifB*-III and the other two parts (*nifB*-I and II), the full-length *nifB* coding frame cannot be restored by simple excision of the insertion sequences. Furthermore, the *nifB* and *nifH1* parts are alternatively encoded: *nifH1*-IV, *nifB*-II, *nifB*-I, *nifH1*-III, *nifB*-III, *nifH1*-I, and *nifH1*-II. For example, the 72-kb region between *nifB*-I and *nifB*-III contains the *nifH1*-III fragment, and the 58-kb region between *nifH1*-III and *nifH1*-I contains the *nifB*-III fragment and the three contiguous genes, *fdxN-nifS-nifU*. Therefore, if the insertion sequence is simply excised to restore either *nifH1* or *nifB*, one of the fragments of *nifB* and the three genes or *nifH1* will be lost from the chromosome. Given this high degree of fragmentation and the inconsistent coding order with opposite directions in some parts of the *nifBH1DK* genes, NIES-4101 may have already lost its nitrogen-fixing activity or could be expressing functional *nif* genes in a heterocyst cell-specific manner through multi-step programmed genome rearrangements that are much more complex than in the those of *A.*7120.

### Heterocyst differentiation and diazotrophic growth

NIES-4101 cells grown under nitrogen-fixing conditions (BG-11_0_) were observed under an optical microscope (Fig. 2A, C). Heterocyst cells were observed at the terminal end, and tapering trichomes were observed toward the opposite end. Fluorescence microscopy (Fig. 2B, D) revealed that only the terminal cells showed extremely low chlorophyll fluorescence, which is characteristics of heterocysts. Such morphological features are consistent with the characteristics of the *Calothrix* genus (Rippka et al. 1979).

**Figure 2.**
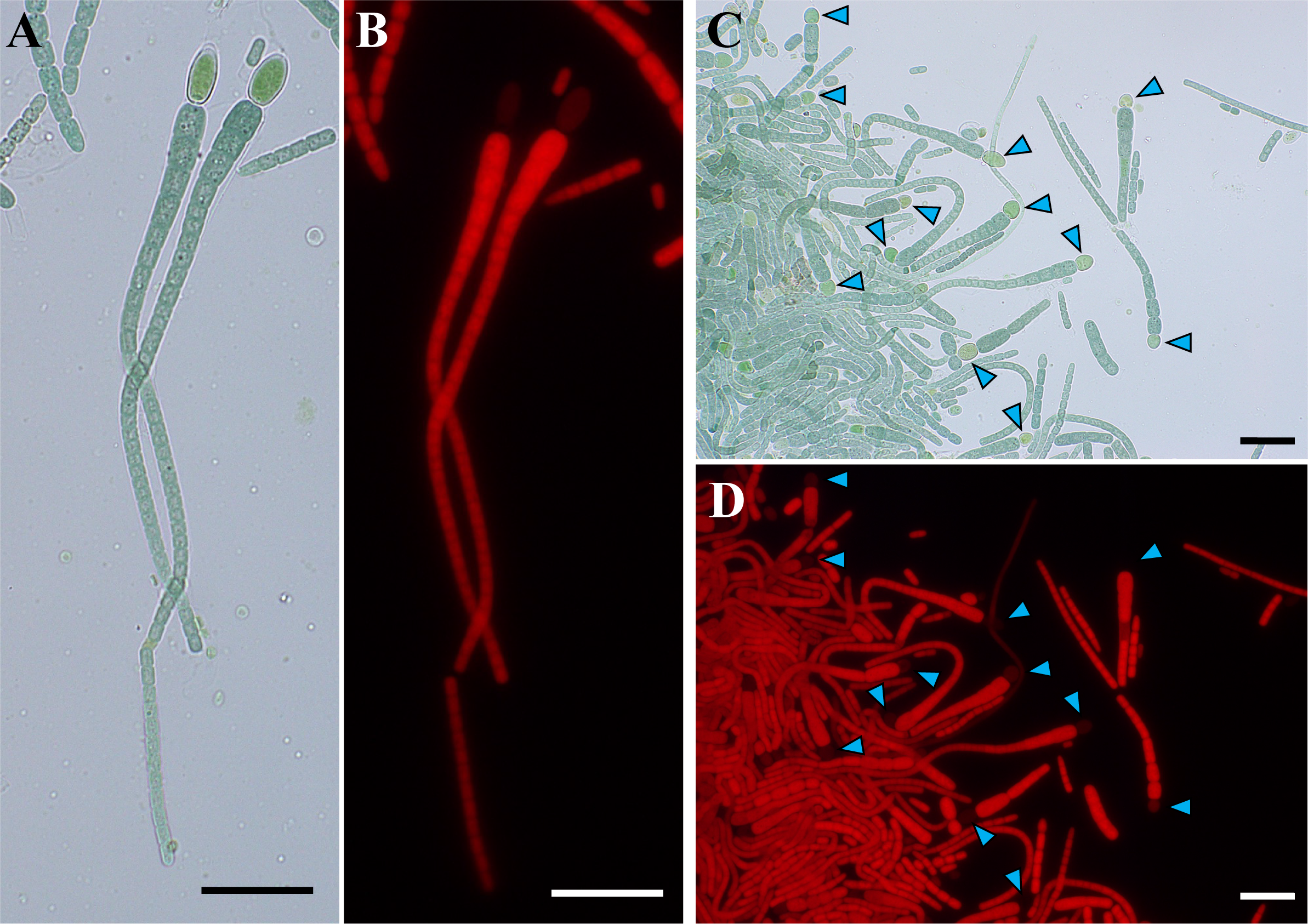
Optical microscopic observation of NIES-4101. Trichomes of NIES-4101 in bright field (**A**, **C**) and in chlorophyll autofluorescence excited at 554 nm (**B**, **D**). Heterocyst cells are indicated with light blue arrowheads. NIES-4101 cells grown on nitrogen-free BG-11_0_ agar plates for 14 days. Scale bars = 20 µm.

The diazotrophic growth and nitrogenase activity of NIES-4101 was examined. NIES-4101 cells grown on a BG-11 agar plate formed tightly aggregated cell clumps, which could not be easily dispersed. A cell clump of approximately 5 mm in size was loosened with tweezers and inoculated onto a BG-11 (15 mM KNO_3_) or a BG-11_0_ (without combined nitrogen) agar slant in a 50-mL glass vial. Cells grew to the same extent in both conditions, strongly suggesting that NIES-4101 grew on the BG-11_0_ slant by nitrogen fixation (Supplementary Fig. S1).

### Nitrogenase activity

The nitrogenase activity of NIES-4101 was examined using acetylene reduction. Due to the difficulty of preparing homogeneous cell suspensions of NIES-4101 as described above, the nitrogenase activity was measured as the acetylene reduction activity for each slant vial containing an almost similar cell clump (Fig. 3). Acetylene reduction activity was measured in the BG-11_0_ slant vials every 7 days after inoculation, using a BG-11 slant vial as a negative control. Significant acetylene reduction activity was detected only in BG-11_0_ slant vials (Fig. 3). In contrast, no acetylene reduction activity was detected in the BG-11 slant vials after 7 and 14 days, and only low activity was observed after 21 and 28 days. The growth on a BG-11_0_ slant was comparable to that on BG-11, and the acetylene reduction activity was detected only in BG-11_0_ slant vials, confirming that NIES-4101 has the capability of growing by nitrogen fixation with nitrogenase.

**Figure 3.**
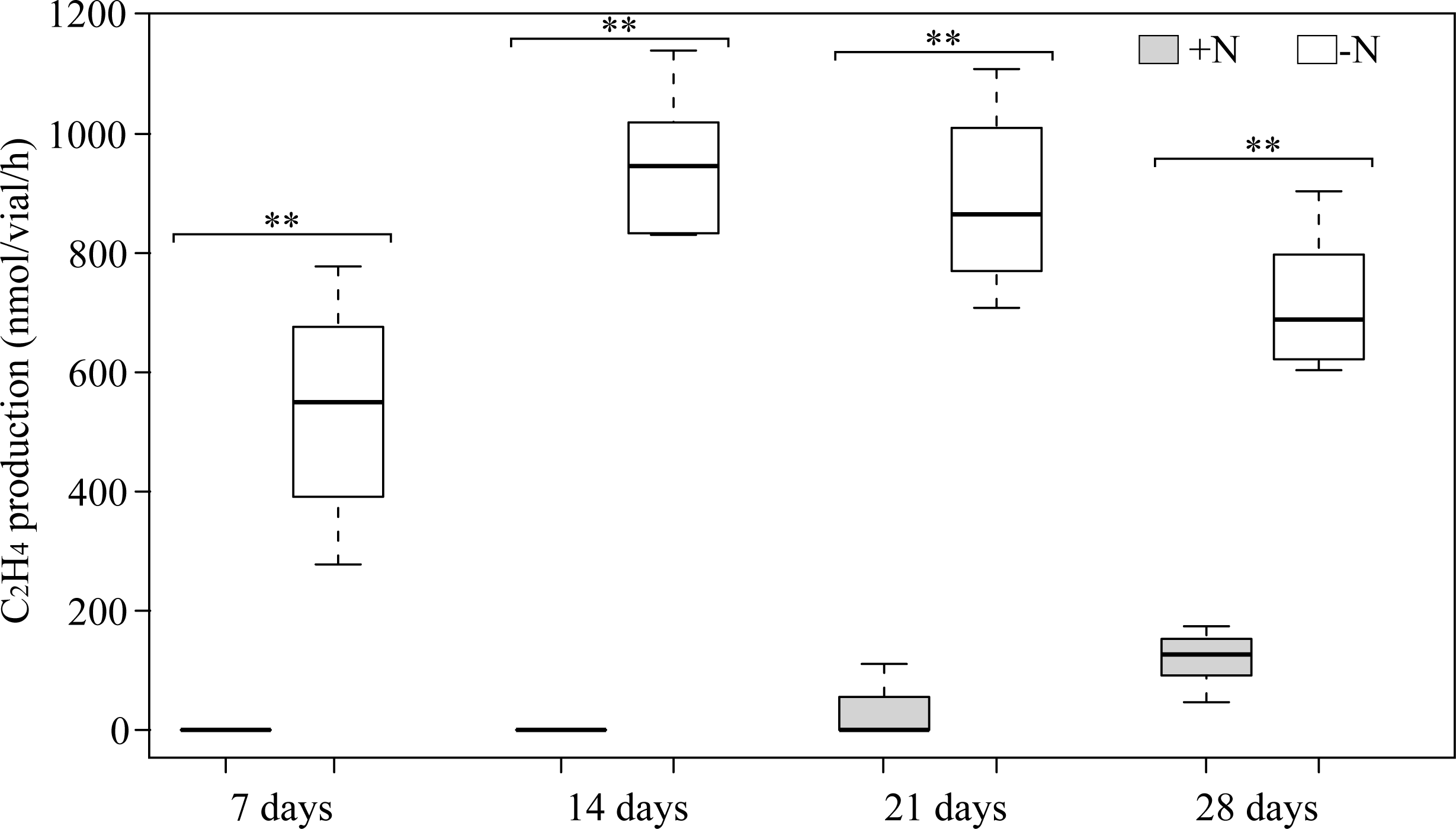
Nitrogenase activity of NIES-4101 estimated by ethylene formation. NIES-4101 cells grown on a BG-11 agar plate were inoculated onto slant vials of BG-11 (gray) or BG-11_0_ (white), followed by incubation under aerobic and light (20 µmol m^-2^ s^-1^) conditions at 30°C. The acetylene reduction activity in each vial was measured (eight for each condition) after 7, 14, 21, and 28 days. The error bar indicates standard deviations. Double asterisk indicates p < 0.001 between BG-11 and BG-11_0_ vials (unpaired t-test). Photographs of the vials are supplied in Supplementary Figure S1.

### Restoration of the functional nif genes: resequencing analysis

The diazotrophic growth and high level of acetylene reduction activity under combined nitrogen-free conditions indicated that NIES-4101 heterocyst cells express a functional nitrogenase, suggesting that the functional *nifBH1DK* genes with the complete coding frames are formed by complex recombination events from the highly fragmented parts in heterocysts. To obtain direct evidence for the genome rearrangement in the *nifBH1DK* genes, genomic DNA was prepared from NIES-4101 cells grown in nitrogen-fixing conditions. A DNA library was prepared, and bulk short-read sequencing was performed. An *in-silico* pipeline (Supplementary Fig. S2) to enrich for paired-end reads sequenced across the possible breakpoints of the rearrangement yielded 154,000 × 2 paired-end reads (representing 3.4% of all shotgun reads). *De novo* assembly of these reads yielded two heterogeneous contigs with 100% sequence identity to the respective fragmented *nifBH1DK* sequence. A 4,879-bp contig contained the full length of the *nifH1*, *nifD*, and *nifK* genes (Fig. 1A). This result clearly indicates that the *nifH1DK* operon is restored by a series of rearrangement events in the heterocysts. For *nifD* and *nifK*, it is assumed that the complete coding frames are restored by simple repeat of excision events. The average rate of reads corresponding to the seven recombination sites to which each part of *nifH1DK* is connected during the restoration process accounted for 4.1% of the total reads which sequenced at each breakpoint.

Another 3,264-bp contig was found for the full-length *nifB* coding sequence. This contig contained the full-length *nifB* coding frame, the 1.1-kb non-coding upstream region of *nifB*, and the *fdxN* gene located just downstream of *nifB* (Fig. 1A). In the genome from the vegetative cells, the three contiguous genes *fdxN-nifS-nifU* were encoded just downstream of *nifB*-III in the same direction. However, this orientation is opposite to that of the *nifDK* parts; thus, the orientation must be reversed to the opposite direction during the restoration process. The direction of the *nifB*-II and *nifB*-I fragments was consistent with that of *nifDK* (Fig. 1A). The average rate of reads corresponding to the two recombination sites for the reconstructed *nifB* gene accounted for 5.5% of all reads which sequenced at each breakpoint. Considering the results from *nifH1DK*, an average of 4.4% of reads were derived from the rearrangement junctions, suggesting that approximately 4%–5% of all NIES-4101 cells differentiate into heterocysts. The restoration process of *nifH1* and *nifB* will be discussed later.

### Reconstitution of the functional nif genes: PCR analysis

To verify the reconstitution of the *nifBH1DK* genes, a PCR experiment was performed with the appropriate primers. Six DNA fragments covering the individual coding regions of *nifB*, *nifH1*, *nifD*, *nifK*, *nifH1-nifD-nifK*, and *fdxN*-*nifS*-*nifU* were amplified (Supplementary Fig. S3). The nucleotide sequences of the amplified fragments were determined using the Sanger method, and the amino acid sequences showed high similarity to those of the orthologs (Supplementary Figs S4–7). We also confirmed the presence of the restored *nifB* operon adjacent to the *nifEN* operon (Fig. 1B). In this PCR reaction, the forward primer, nifE_F1, corresponding to the *nifE* coding region and the reverse primers, nifB_F2 and nifBpro_R1, corresponding to near the 5L-region of *nifB* and to the *nifB* promoter, respectively, were used. In the genome of vegetative cells, the size of the estimated PCR fragment by the nifE_F1/nifBpro_R1 primer combination was approximately 260 kb, which is too long to amplify by normal PCR. In the nifE_F1/nifB_F2 primer combination, amplification did not occur because nifB_F2 is orientated in the same direction as nifE_F1 in the genome of vegetative cells. If the functional *nif* operon is formed in the genome of heterocyst cells, the primer pairs, nifE_F1/nifB_F2 and nifE_F1/nifBpro_R1, allow the amplification of 8,449-bp and 10,537-bp fragments, respectively, spanning *nifB-fdxN-nifS-nifU-nifH1-nifD-nifK-nifE*. Figure 1B shows the successful amplification of the 8.4-kb and 10.5-kb fragments with primers nifE_F1/nifB_F2 and nifE_F1/nifBpro_R1, respectively. Detection of the two contigs reconstructed in *de novo* assembly from resequencing reads and the PCR results strongly suggest that the 13 fragmented parts of the *nifBH1DK* genes scattered throughout the 350-kb region are correctly restored to form the *nifB* operon next to the adjacent the *nifEN* operon, resulting in the 17-kb *nif* gene cluster. It should be noted that the gene arrangement of the reconstituted *nif* operon in NIES-4101 is well conserved in various heterocyst-forming cyanobacteria, including *A.* 7120 and *Trichomonas variabilis* ATCC 29413 (Supplementary Fig. S8) (Böhme 1998; Thiel 2019). In addition, it should be noted that in the promoter region immediately upstream of *nifB* there is the *cis* element that is recognized by CnfR, the master transcriptional regulator in nitrogen fixing cyanobacteria (Supplementary Fig. S9) (Tsujimoto et al. 2016).

## Discussion

### Overview

In this study, we found that in the genome of NIES-4101, the four *nifBH1DK* genes, which are essential for nitrogen fixation, were fragmented into 13 parts scattered across a 350-kb chromosomal region. We identified contigs and PCR fragments covering the fully restored *nifBH1DK* genes. The restored *nifBH1DK* subcluster is linked to the *nifEN* subcluster, then completing the 17-kb *nif* gene cluster. This experimental observation is consistent with the nitrogen-fixing growth and the nitrogenase activity in NIES-4101. The highly fragmented *nif* genes are restored in the full-length *nifBH1DK* genes by a complex and precise recombination mechanism.

### A model for restoration of the nif gene cluster

We proposed a model for the genome rearrangement of the *nif* cluster (Fig. 4). Since the *nifK* and *nifD* fragments are encoded in the same direction, the full-length coding frames are restored by simple excision events that remove the insertion sequences, which are catalyzed by site-specific recombinases that are encoded in the insertion sequences (Fig. 4A). This type of excision is similar to that of the 11-kb insertion sequence by XisA in the *A.* 7120 *nifD* gene (Golden et al. 1985).

**Figure 4.**
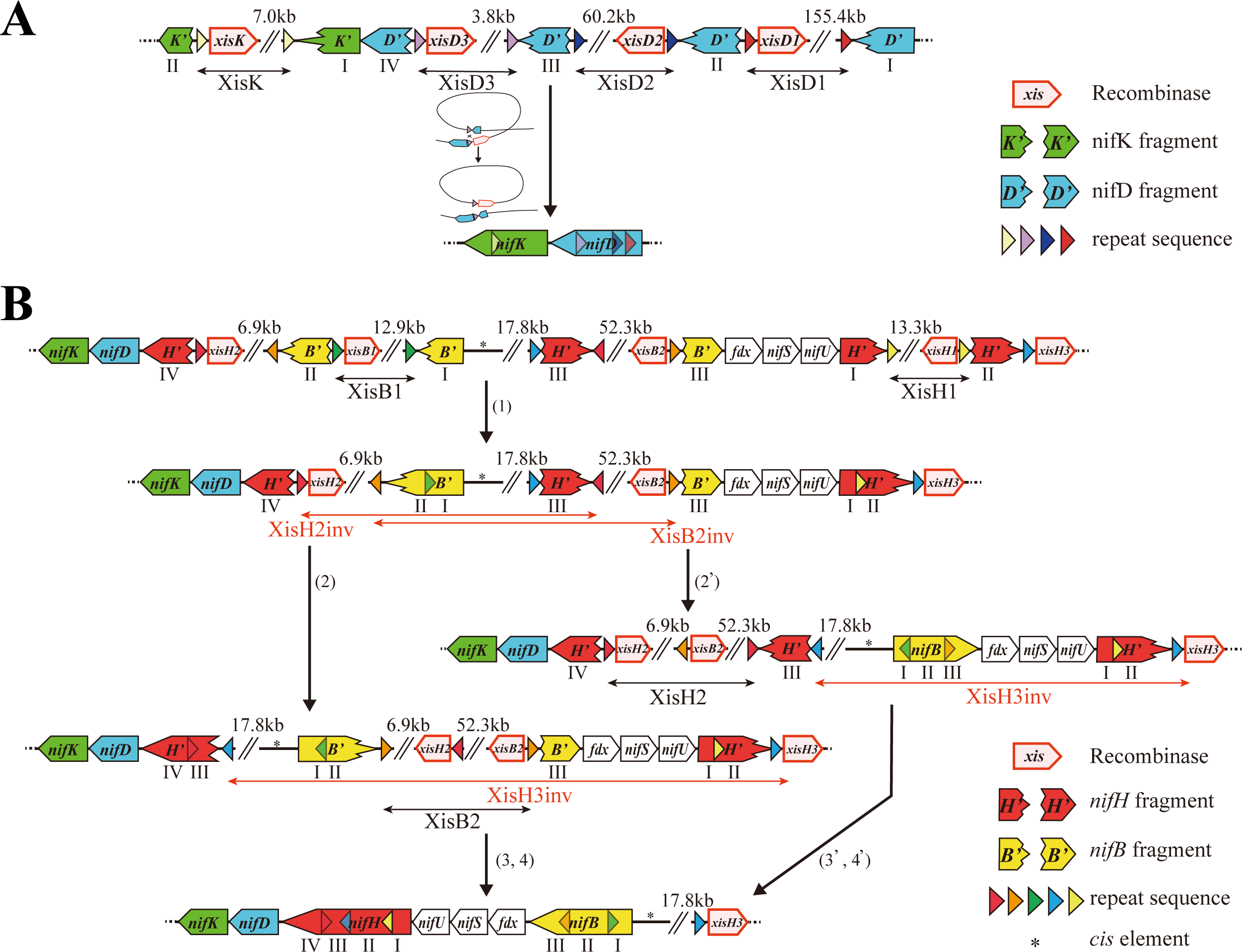
A model for the restoration of the *nif* gene cluster. (**A**) Restoration of *nifK* and *nifD*. The full *nifD* and *nifK* genes are reconstituted by four simple excision events. (**B**) Restoration of the *nifH1* and *nifB*. (1) The sequences between the direct repeats (black bars with arrowheads at both ends) are excised by the site-specific recombinases encoded adjacent to the respective repeats. (2) Inversion occurs at the inverted repeats (red triangles, red bar with arrowheads at both ends) between the *nifH1* fragments (*nifH1*-III) and (*nifH1*-IV). (2’) Alternatively, inversions occur at the inverted repeats (orange triangles, red bar with arrowheads at both ends) adjacent to the *nifB* fragments (*nifB*-I/II) and (*nifB*-III). (3) Inversion occurs at the inverted repeats (blue triangles, red bar with arrows at both ends). (4) The insertion sequence between the *nifB* fragments (brown triangles, black bar with arrowheads at both ends) is excised to form the full-length *nifB* gene. Alternatively, (3’) inversion occurs at the inverted repeats (blue triangles, black bar with arrows at both ends). (4’) The insertion sequence between the *nifH1* fragment is excised (red triangles, black bars with arrowheads at both ends) to form the full-length *nifH1* gene. Genes in the process of restoration are indicated with primes (*H’* and *B’*). The unrelated region is indicated by “//”. Recombinase gene names are shown without the prefix Cn, which indicates the species (*<u>C</u>alothrix* sp. <u>N</u>IES-4101). Recombinases that catalyze inversions (CnXisH2, CnXisB2, and CnXisH3) are indicated with “inv” at the end of the name.

However, to restore the complete *nifH1* and *nifB* genes, change of the direction via an inversion event is needed for some fragments. Considering that recombinase has the activities to excise the region surrounded by the direct repeat sequences and to invert the region surrounded by the inverted repeat sequences (Abremski et al. 1983), we propose a complex recombination mechanism that includes excision and inversion events as follows (Fig. 4B):

1. Recombination events between the direct repeat sequences of *nifH1*-I and *nifH1*-II (Fig. 4B, yellow triangles) and between *nifB*-II and *nifB*-I (Fig. 4B, green triangles) excise the insertion sequences resulting in the formation of the intermediate fragments, *nifH1*-I/II and *nifB*-I/II, respectively.
2. Recombination events between the inverted repeat sequences next to *nifH1*-IV and *nifH1*-III (Fig. 4B, red triangles) invert the 27.5-kb region to form another intermediate fragment, *nifH1*-III/IV.
3. A recombination event between the inverted repeat sequences of *nifH1*-III and *nifH1*-II (Fig. 4B, blue triangles) inverts the 100-kb region to restore the full *nifH1* gene.
4. A recombination event between the direct repeat sequences of *nifB*-II and *nifB*-III (Fig. 4B, orange triangles) excises the insertion sequence (62.4 kb) to restore the full *nifB* gene.

Alternatively, after step (1),

(2’) A recombination event between the inverted repeat sequences next to *nifB*-II and *nifB*-III (Fig. 4B, orange triangles) inverts of the 73.2-kb region to restore the full *nifB* gene.

(3’) A recombination event between the inverted repeat sequence of *nifH1*-III and *nifH1*-II (Fig. 4B, blue triangles) inverts of the 23.0-kb region to form an intermediate connecting fragment *nifH1*-I/II/III.

(4’) A recombination event between the direct repeat sequences (Fig. 4B, red triangles) of *nifH1*-IV and *nifH1*-III excises the insertion sequence (62.4 kb), resulting in the restoration of the full *nifH1* gene.

Upon completion of the sequential rearrangement events (1)–(4) or (1)–(4’), the functional *nifB-fdxN-nifS-nifU-nifH1-nifD-nifK* cluster is eventually restored (Fig. 4B, bottom). Then, immediately downstream of this *nifB* subcluster, there is another subcluster of *nif* genes; *nifE-nifN-nifX*, *nifW*, *hesA*, and *hesB*-*fdxH*, which completes the functional 17-kb *nif* gene cluster (Supplementary Fig. S8).

### Site-specific recombinases

In the proposed model, nine recombination events occur to reconstitute the functional *nif* gene cluster. The genes encoding the recombinases that are responsible for the recombination events are found in the vicinity of the fragmented parts of the *nifBH1DK* genes, most of which are located within the insertion sequences to be excised or inverted (Figs. 1A, 4; Table 2). We hypothesize that seven of them catalyze the excision of the insertion sequences, and the other two catalyze the inversion of the regions flanked by the inverted repeat sequences. Based on the gene name and order of the target genes linked by the excision or inversion event, we propose the following designations for these nine recombinases (Figs. 1A, 4; Table 2): CnXisD1, CnXisD2, and CnXisD3 for the *nifD* recombinases, starting from the N-terminus of NifD; CnXisK for the *nifK* recombinase; CnXisB1 and CnXisH1 for the first recombinases acting for *nifB* (*nifB*-I/II) and *nifH1* (*nifH1*-I/II), respectively; CnXisH2 and CmXisH3 for the two successive inversion events, (2) and (3), respectively; and CnXisB2 for the final recombinase to restore the full *nifB* gene. For the alternative route, CnXisB2 and CnXisH3 could undergo another two successive inversion events, (2’) and (3’), respectively, and CnXisH2 for the final recombinase to restore the full *nifH1* gene.

**Table 2.**
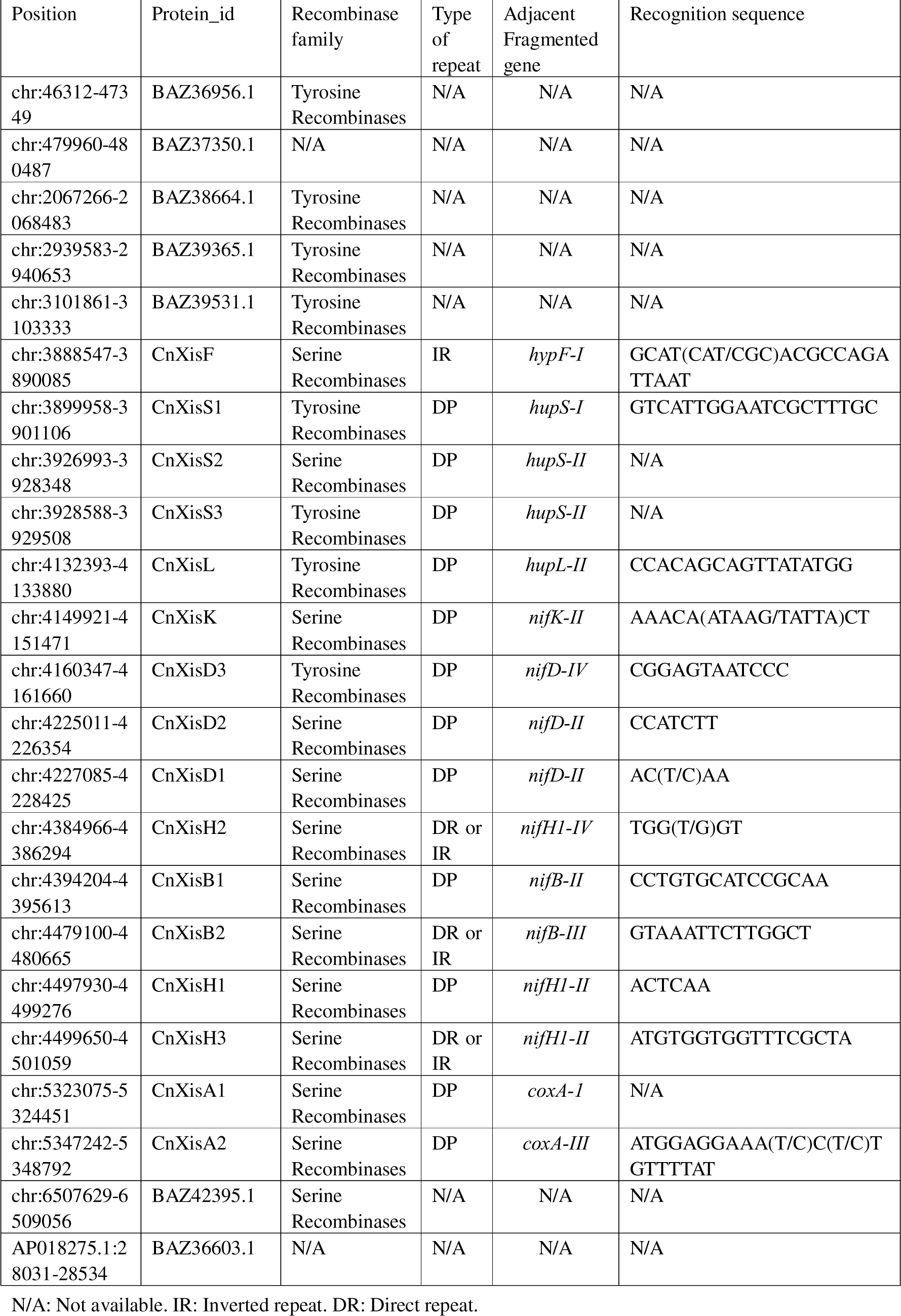
List of putative site-specific recombinases in NIES-4101.

In this model, the second inversion reaction by CnXisH3 ((3) or (3’)) cannot occur without the first inversion reaction by CnXisH2 (2) or CnXisB2 (2’). In addition, the direct repeat sequence to join the *nifB* (4) or *nifH1* (4’) fragments cannot occur without the first inversion reaction. Therefore, the series of recombination reactions must occur in sequential order, with the inversion reaction by CnXisH2 (2) or CnXisB2 (2’) as the first inversion reaction, and then by CnXisH3. After the first inversion reaction, the *nif* cluster restoration can be completed even if either the inversion reaction by CnXisH3 ((3) or (3’)) or the excision reaction by CnXisB2 (4) or CnXisH2 (4’) occurs first. In other words, the two inversion events change the direction of the *nifH1* and *nifB* fragments in the opposite strand, allowing for the final reconstruction of *nifH1* and *nifB* by simple excision events.

When all recombination events are completed, the eight genes for site-specific recombinases are removed from the chromosome, while only one gene for CnXisH3 remains 90-bp downstream of the repeat sequence (Fig. 4, blue triangle). If CnXisH3 is still expressed in heterocysts after the completion of the reconstruction of the *nif* gene cluster, it is possible that the second inversion reaction ((3) or (3’)) is in equilibrium in heterocyst cells. This situation is difficult to verify by PCR or short-read genome resequencing because the sequence before the inversion reaction is the identical to that of the chromosome of vegetative cells. The ratio of sequencing reads for the *nifH*-II and *nifH*-III junctions to the total reads was 5:238 (2.1%), which is close to that of the background reads and not so significantly different from that of other junction sites (Supplementary Table S1). This result is not inconsistent with the status where almost all chromosomal copies are restored in the heterocyst chromosome.

Recombinases that catalyze site-specific recombination are divided into two groups: tyrosine recombinases and serine recombinases (Grindley et al. 2006). The sequence similarity of the recombinases indicated that, except for CnXisD3, which is classified as a tyrosine recombinase, all others are serine recombinases (Table 2). CnXisD3 showed the highest similarity (81.5% identity) to XisA of *A.* 7120 (Lammers et al. 1986) (Supplementary Fig. S12, Supplementary Table S2). The high sequence similarity between CnXisD3 and XisA is reflected in their recognition sequences: the direct repeat sequences recognized by CnXisD3 and XisA are GGGATTACTCCG (12 bp) and GGATTACTCCG (11 bp; Golden et al. 1985), respectively (Table 2), which are nearly identical. The excision of the insertion sequence at the 3’ end of the *nifD* gene is highly conserved in NIES-4101 and *A.* 7120 in terms of similarity of the recombinase, the local amino acid sequences corresponding to the excision site, and the recognition sequence by the recombinases, suggesting a common evolutionary origin.

The other eight recombinases were classified as serine recombinases, similar to XisF in *A.* 7120 (Table 2) (Carrasco et al. 1994). Two accessory proteins, XisH and XisI, encoded in the tight operon *xisH-xisI*, are reported to be required for the XisF activity (Ramaswamy et al. 1997). In the genome of NIES-4101, there are three *xisH-xisI*-like operons encode proteins that show significant similarity to XisH and XisI (chr: 155,515–156,257 (NIES4101_29730 and NIES4101_49740), 4,035,268–4,036,007 (NIES4101 and 62630-NIES 4101_62640) and 6,800,108–6,800,856 (NIES4101_86010 and NIES4101_86020)). Since none of the three operons are present in the insertion sequences of the *nif* region, it is not clear whether these proteins are involved in the eight recombinases, nor are they specific to some recombinases. Of the eight recombinases, CnXisK and CnXisB2 showed the highest amino acid sequence similarity to XisF (49.8% and 39.1%, respectively) (Supplementary Table S2, Fig. S12).

The seven site-specific excision reactions in the reconstitution process of the *nif* gene cluster in NIES-4101 produce double-stranded circular DNAs in which the insertion sequence is circularized. The 11-kb insertion sequence of *nifD* has been detected as double-stranded circular DNA in *A.* 7120 (Golden et al. 1985). In NIES-4101, contigs with the ends of the insertion sequences connected to each other were detected in the resequencing reads, suggesting that the excised insertion sequences in the seven loci remain as double-stranded circular DNA in the heterocyst cells.

### Activities of recombinases

In the genome of vegetative cells, there are two pairs of inverted repeats are recognized by the recombinases CnXisH2 and CnXisB2, and which of the two pathways ((2)–(4) and (2’)–(4’)) is selected depends on whether CnXisH2 or CnXisB2 acts first. When CnXisH2 acts first, the pathway (2)–(4) is selected, and the restoration is completed by the inversion by CnXisH3 and the excision by CnXisB2. When CnXisB2 acts first, it follows the pathway (2’)–(4’), and the restoration is completed by the inversion by CnXisB2 and the excision by CnXisH2. It is important to study in future work how the order of action of recombinases is determined.

### Another model for reconstitution via double-stranded circular DNA

We also hypothesize an alternative model without any inversion events (Supplementary Fig. S10). After the two excision events in the previous model (1), an excision event occurs between the direct repeats flanking *nifH1*-III and *nifH1*-II (blue triangles). Then a region of approximately 58.6-kb is excised as double-stranded circular DNA (an episome), in which an intermediate fragment, *nifH1*-I/II/III, is formed. When this circular DNA element again undergoes recombination between the direct repeats (red triangles) next to *nifH1*-III (circular DNA element) and *nifH1*-IV (chromosome), the circular DNA is reincorporated into the chromosome, resulting in the formation of the full *nifH1*, followed by a single simple excision event (orange triangles) to form *nifB*, giving rise to the full *nifBH1DK* genes. This episome-mediated reconstitution can be completed with a simple repetition of excision events between direct repeat sequences. It consists of five recombination events, two of which are the excision events to produce the episome and the re-introduction event of the episome into the chromosomes. The total number of recombination events, five, is the same as that in the first model, including two inversion events (Fig. 3B). Further studies, including assays of recombinase activity and detection of the episome, are needed to establish which recombination mechanisms occur during the heterocyst development.

### Excision events of insertion sequences other than the nif genes

We have preliminary obtained results indicating the reconstitution events that occur in the four other genes: *hupL*, *hupS*, *hypF*, and *coxA* genes (Supplementary Fig. S11). The full *hupL* gene is restored by a simple single excision event of the 40-kb insertion sequence. However, the *hupS* and *coxA* genes are fragmented into three parts, which need two excision events to restore the full genes. The *hypF* gene is fragmented into two parts, one of which is encoded in the reverse strand, which require a single inversion event to restore the full gene. Seven recombinases appear to involve the site-specific excision and inversion events (Supplementary Table S1, Fig. S12). However, at this time, only four recombinases (*hupL*, *hupS* (I/II), *hypF*, and *coxA* (II/III)) are assumed their recognition repeat sequences (Table 2, Supplementary Table S1). Interestingly, the recognition sequence (17-bp; CCACAGCAGTTATATATGG) for the *hupL* excision is nearly identical to that in *A.* 7120 for the *hupL* excision (15-bp; ACAGCAGTTATATATGG; Carrasco et al. 1995), and the recombinases of CnXisL and XisC show very high similarity (86%), suggesting that the *hupL* excision, similar to the *nifD* excision, is evolutionarily conserved. For the other three, no appropriate repeat sequences have been found. In the NIES-4101 genome there are 26 genes encoding recombinases (Supplementary Table S1). Therefore, it is still possible to find other fragmented genes in the genome.

### Distribution of fragmented nif genes in the Calothrix genus and related genera

Prevalence of fragmentation of *nif* genes has been reported previously in heterocyst-forming cyanobacteria (Hilton et al. 2016), in which the authors regarded some fragments encoded in the reverse strand as misassemble of genome sequencing data. To investigate whether the fragmentation of *nif* genes and the restoration system including inversion are evolutionarily conserved in the *Calothrix* genus and other closely related genera, we examined genomes from 13 *Calothrix* strains, 7 *Fischerella* strains, 4 *Rivularia* strains and other closely related 4 strains (Fig. 5, Supplementary Table S3; total 29 including NIES-4101 with two strains *T. variabilis* ATCC 29413 and *A.* 7120 as outgroups) available at NCBI assembly based on the Genome Taxonomy Database (GTDB) classification (Parks et al. 2020). In particular, we focused on the six core *nif* genes, *nifHDKENB*, and investigated the number of fragments and their direction (Fig. 5, Supplementary Table S4). Regarding the number of fragments, no fragmentation was found in 5 of the 29 species. Four of them were in the genus *Fischerella* (three of which were reported in Hilton et al. (2016)), and only one strain, *Calothrix* sp. 336/3, in the *Calothrix* genus, showed no fragmentation. In the other 24 strains, fragmentation in at least one *nif* gene was observed. Except for five strains in which only one gene is fragmented into two parts (*nifD*, three strains; *nifK*, one strain; and *nifH*, one strain), the other 19 strains showed fragmentation in two or more *nif* genes. Interestingly, no strains with only fragmented *nifENB*, and fragmentation tended to occur mainly in *nifHDK*. In addition, the number of fragmentations tends to be more in the genera *Rivularia* and *Calothrix* than in *Fischrella*.

**Figure 5.**
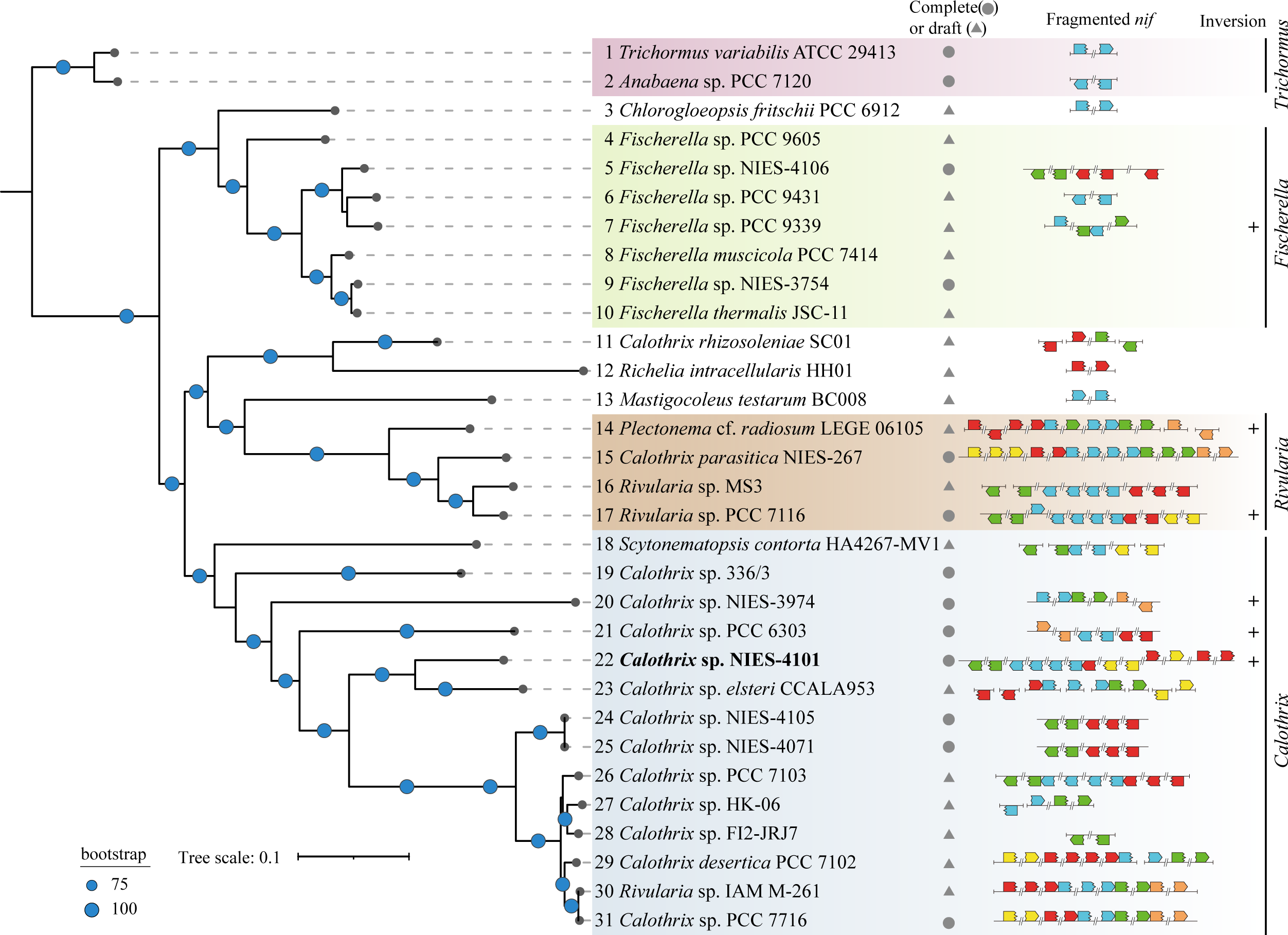
Core protein phylogeny of NIES-4101 and closely related cyanobacterial strains. The 59 core gene-set was estimated using Roary v3.13.0 from genome assemblies of 14 *Calothrix* strains, 7 *Fischerella* strains, 4 *Rivularia* strains, and 2 *Trichormus* strains, and 4 closely related strains (*Chlorogloeopsis fritschii* PCC 6912, *Calothrix rhizosoleniae* SC01, *Richelia intracellularis* HH01, and *Mastigocoleus testarum* BC008). *T. variabilis* ATCC 29413 and *A.* 7120 were used as outgroups. Tree inference was performed using the best-fit model in IQ-TREE v2.2.0.3 with ultra-fast bootstrap 1,000 approximation. Genomes with fragmented *nifHDKENB* genes are indicated by horizontal arrows indicating fragmented *nif* genes. The direction and position of the arrows (red, *nifH*; blue, *nifD*; green, *nifK*; yellow, *nifB*; and orange, *nifE*) indicate the position and orientation of the fragmented *nifHDKENB* genes on the genome assembly. Blank indicates the presence of full-length *nif* genes without insertions. Note that no *nifN* with insertions were found in these strains. A + mark on the right raw indicates at least one fragment is in the opposite direction to the other fragments. Background color for major 4 clade represents genus-level taxonomic classification based on GTDB classification. The phylogenetic tree was constructed using the iTOL v6 web service.

For the arrangement of fragments and the direction, it should be noted that only partial information is obtained for the 18 strains whose genome information is at a draft level, because the distance between fragments (the insertion sequence) may range from a minimum of 1 kb to a maximum of 1 Mb (Hilton et al. 2016) and the orientation of fragments can only be determined when the fragments are encoded in the same contig. We found that six strains including NIES-4101 out of the 24 strains have *nif* gene fragments in the opposite direction (at least one contig for two strains with draft genomes; Fig. 5, inversion +). *nifD* and *nifK* in *Fischerella* sp. PCC 9339 were fragmented into two parts and each one part is in the opposite direction in one contig. In this case, we can easily assume that the complete *nifDK* operon is restored by inversion of the fragment containing the contiguous *nifDK* parts in the reverse strand. In two strains of the *Rivularia* genus, *Plectonema* cf. *radiosum* LEGE 06105 and *Rivularia* sp. PCC 7116, only ones of the four fragments of *nifH* and of the five fragments of *nifD* are in the opposite direction, respectively. In two strains of *Calothrix* sp. NIES-3974 and PCC 6303 both have only one of the two *nifE* fragments was in the opposite direction. These results suggest that the restoration system involving inversion is not limited to NIES-4101, but is distributed scattered throughout the genera *Fischerella* and *Calothrix*. Of the 29 strains compared in this study, we found that NIES-4101 has the most complex restoration system consisting of nine recombination events of excision and inversion.

### Recombinases catalyzing inversion

We examined whether the recombinases catalyzing inversion are evolutionarily related. We constructed phylogenetic trees of 21 recombinases from NIES-4101 including CnXisH3, which is assumed to catalyze inversion (Fig. 4B), and five recombinases with probable inversion activity from the five strains (Supplementary Figure S12). Interestingly, the recombinase WP_008231949.1.1 from *Plectonema* cf. *radiosum* LEGE 06105, is the most closely related to CnXisH3 with a high similarity of 83%, supporting the idea in which these two recombinases catalyze inversion reaction in common. On the other hand, the other four probable recombinases for inversion are not closely related to CnXisH3, but rather to a variety of other recombinases catalyzing excision such as CnXisD2 and CnXisD3. This phylogeny suggests that the recombinases catalyzing inversion acquired the inversion activity independently during evolution.

### Paired-end sequencing technology

In this study, we used a paired-end sequencing technology to reconstruct the *nif* gene cluster; single DNA molecules of 300–1,000 bases in length were sequenced from both sides. Paired-end reads sequenced spanning the recombination breakpoint were collected and assembled *de novo*. This allowed us to assemble heterogeneous contigs within a few hundred base pairs spanning the rearrangement breakpoints. Since the *nifH1*, *nifD*, and *nifB* genes of NIES-4101 are interrupted by multiple insertion sequences, it is even more advantageous for full-length reconstitution. The simple *in silico* approach proposed here is expected to enable low-cost reconstitution of genes where complex recombination has occurred.

## Materials and Methods

### Culture conditions

*Calothrix* sp. NIES-4101 was provided from the NIES collection (Microbial Culture Collection at the National Institute for Environmental Studies, Tsukuba) in December 2022. BG-11 and BG-11_0_ (Rippka et al. 1979, Tsujimoto et al. 2014) agar plates adjusted to pH 7.5 with HEPES buffer (20 mM) were used for cultivating NIES-4101 in nitrogen-sufficient and nitrogen-fixing conditions, respectively. Agar plates were incubated aerobically at 30°C under continuous light conditions (20 µmol m^−2^ s^−1^). NIES-4101 cells grow as tightly aggregated colonies on agar plates and cannot be easily dispersed. When inoculating onto a new agar plate, cell aggregates were loosened with tweezers and a cell aggregate of approximately 7-mm^2^ size was transferred onto a new agar plate.

### Acetylene reduction activity

NIES-4101 cells cannot be suspended homogeneously in a liquid medium due to them forming tightly aggregated colonies. The nitrogenase activity of NIES-4101 was measured as acetylene reduction activity for each 50-mL vial containing agar slant medium (13 mL). Agar slants of BG-11 or BG-11_0_ medium (13 mL) prepared in a 50-mL vials (Cat No. 0501-08, Nippon Electric Glass, Otsu) were inoculated with a cell aggregates (approximately 7 mm^2^) of NIES-4101 grown on a BG-11 agar medium. The 50-mL vial was covered with an air permeable silicone cap and incubated under continuous light conditions (20 µmol m^−2^ s^−1^). The nitrogenase activity in each vial was measured every week (Fig. 3). During activity measurement, the air-permeable silicone cap was replaced with an air-tight silicon septum, and the gas phase of the vial was replaced by sparging with an acetylene gas mixture (10% acetylene-90% argon) for 15 sec. The vial was incubated at 30°C under continuous light conditions for 30 min. A portion of the gas phase (500 µL) was analyzed by gas chromatography as described previously (Tsujimoto et al. 2014). After the measurements, the cap of the vial was replaced with an air-permeable silicone cap and the incubation was continued (Supplementary Fig. S1).

### Preparation of genomic DNA

NIES-4101 cells grown on a BG-11 agar plate were transferred onto a piece (5 cm^2^) of sterile Hybond N+ membrane placed on a BG-11_0_ agar plate to grow diazotrophically for 2 weeks at 30°C under continuous light conditions (20 µmol m^−2^ s^−1^). Cell aggregates grown on the membrane were transferred to a 2-mL microtube and prewashed once with sorbitol solution containing 1% (v/v) β-mercaptoethanol to remove secondary metabolites. After centrifugation at 7,000 rpm (rotor, RT15A3; model, himac CF 15D2; Eppendorf Himac Technologies, Hitachinaka), the supernatant was discarded. Then, 1 mL of 1 × TE (10 mM Tris-HCl; pH 8.0, 1 mM EDTA) was added and the sample was frozen at –80°C. After transferring the frozen sample to a mortar, the sample was ground for several minutes with a pestle in liquid nitrogen. The powder was frozen again at –80°C. After another freeze-thaw cycle, a 0.5-mL aliquot of the cell suspension was used for subsequent genomic DNA extraction.

Lysozyme (final 10 mg mL^−1^) was added to the suspension and the sample was incubated at 37°C for 1 h. The suspension was centrifuged once (6,000 rpm for 1 min; RT15A3), and the precipitate was suspended in 100 µL of RNaseA (DNase free, final 20 µg mL^−1^) containing TE buffer. Then, 600 µL of DNA Suisui F (Rizo, Tsukuba) was added (Tomatsu et al. 2018), the sample was mixed well by inversion, and then the sample was incubated at 70°C for 10 min for cell lysis. The cell lysate was transferred to a 2-mL microtube, 800 µL of TE-saturated phenol was added, and the sample was gently stirred with a reciprocal shaker (model, NR-1; TAITEC, Koshigaya) for 15 min. After centrifugation (15,000 rpm for 10 min; RT15A3), the upper water phase was transferred to a new 2-mL microtube. The same procedure was repeated with phenol-chloroform-isoamyl alcohol and chloroform. Genomic DNA was precipitated by adding a 1/10 volume of sodium acetate (3 M, pH 5.2) and the same volume of 2-propanol. The precipitate was washed with 70% ethanol, air-dried, and dissolved in 1 × TE. After the lysis step, all procedures with a micropipette were performed using a wide bore tip.

### Whole-genome sequencing

Short-read sequencing was performed by Genome-Lead (Takamatsu). In brief, DNA extracted for nanopore sequencing (see *Preparation of genomic DNA*) was repurified using AMpure XP SPRI technology (Beckman Coulter, Tokyo). The DNA was then fragmented and size selected using the MGIEasy FS DNA Library Prep Set (MGI Tech Japan, Tokyo), and adapters were ligated. Twelve-cycle PCR amplification was carried out. DNA was heat denatured and single-stranded DNA was prepared. Multiplexed DNA nanoballs were loaded into a flow cell and sequenced on a DNBSEQ-G400RS sequencer (MGI Tech Japan) with 2 × 150 bp chemistry.

### PCR and Sanger sequencing

For PCR reactions, genomic DNA purified with a FavorPrep Genomic DNA Clean-Up Kit (FAVORGEN, Ping Tung, Taiwan) was used as a template. All PCR reactions were performed using PrimeSTAR Max DNA Polymerase (Takara, Kusatsu) with primer pairs (Hokkaido System Science, Sapporo) under conditions that specifically amplify the restored *nif* genes from the genomic DNA of NIES-4101. Sanger sequencing was carried out using an ABI PRISM 3500xL Genetic Analyzer (Sequencing Service, Institute for Gene Research, Nagoya University). The list of primers used for PCR and Sanger sequencing primer is shown in Supplementary Tables S5 and S6, respectively.

### Fragmented nifHDKENB genes

To search for fragmented *nifHDKENB* genes, the NIES-4101 (GenBank accession: GCA_004296455.1) assembly (Hirose et al. 2021) was six-frame searched using tblastn program. The *A.* 7120 NifHDKENB proteins were used as the query sequences, respectively.

### Search for contigs covering the restored nif cluster

In order to assemble the genomic regions where rearrangement occurred in heterocysts, we collected sequencing reads that had been aligned with a partially clipped region against the NIES-4101 reference assembly. The collected paired-end sequencing reads were *de novo* assembled to construct the restored genomic region. The detailed workflow is shown in Supplementary Figure S2. In brief, low quality regions of sequencing reads were trimmed using the fastp preprocessor (-t 1 -T 1 -l 20 -q 20 -n 10 option used)(Chen et al. 2018). The trimmed reads were mapped to the NIES-4101 reference assembly (GenBank accession: GCA_004296455.1) using Minimap2 (Li 2018) and output in SAM format (Li et al. 2009). Alignments containing S (soft clipping) or H (hard clipping) in the CIGAR string were retrieved using awk and grep one-liner. The resynchronized paired-end reads were then *de novo* assembled using MetaSPAdes with the -k 21,31,41,51,61,71,81,91,101,111,121,127 option (Nurk et al. 2017). ORFs encoding Nif proteins were identified using the tblastn program (Gertz et al. 2006) with *A*. 7120 NifHDKENB protein sequences. Two contigs encoding *nifHDK* and *nifB* were manually extended along the genome sequence to reconstruct the full-length *nif* gene cluster. Prodigal (Hyatt et al. 2010) was used to predict the ORF in the *nif* cluster sequence. Since short sequences are insufficient for training *ab initio* gene prediction programs (Trimble et al. 2012), the NIES-4101 genome sequence was temporarily added to the gene prediction program.

### Putative recombinases

Recombinases of NIES-4101 was searched using the blastp program v2.14.1+ (<1e^−20^; Gertz et al. 2006) with the *A.* 7120 XisA, XisC, and XisF as the query sequences. The search was iteratively performed using the found sequences as queries (blastp, <1e^−5^). The recombinases sequences obtained from these operations were reciprocal blastp searched against Siwss-Prot and PDB database (<1e^−5^). Sequences that showed global sequence identity with existing serine or tyrosine family recombinases were considered putative recombinases.

### Phylogenic inference

Thirty-one genome assemblies including 14 *Calothrix* strains, 7 *Fischerella* strains, 4 *Rivularia* strains, and 2 *Trichormus* strains, *Chlorogloeopsis fritschii* PCC 6912, *Calothrix rhizosoleniae* SC01, *Richelia intracellularis* HH01, and *Mastigocoleus testarum* BC008, were downloaded from the NCBI assembly (Supplementary Table S3). Gene prediction was performed using Prokka v1.14.6 (Seemann 2014) with default settings. Core genes were estimated using Roary v3.13.0 (Page et al. 2015) by modifying the protein identity threshold to 90%. Core genes are defined as those present in at least 95% of the strains used. Tree inference was performed using the best-fit model in IQ-TREE v2.2.0.3 (Minh et al. 2020) with ultra-fast bootstrap 1,000 approximation. Genus level phylogenic classification for each assembly was established using GTDB-Tk v2.3.2 (Genome Taxonomy Database Toolkit) with the R207 database (Chaumeil et al. 2022).

## Data Availability

The raw sequencing reads were deposited in the DNA Data Bank of Japan (DDBJ) Sequence Read Archive (SRA) under the BioProject accession number PRJDB16824, and BioSample accession number SAMD00651075, and SRA accession number DRR508037. The reconstituted *nif* gene cluster sequence and annotation were deposited in the DDBJ Nucleotide Sequence Submission System (NSSS) under the accession number LC785442. Additional data, including Sanger sequencing reads, contig sequences, and the commands used to enrich the sequencing reads, are available at GitHub: https://github.com/kazumaxneo/RHGR/tree/main.

## Acknowledgments

We thank Asako Segawa for technical help and Haruki Yamamoto, Takafumi Yamashino, and all members of the Laboratory of Molecular and Functional Genomics for discussion and technical help.

## Author contributions

K.U. and Y.F. conceived the study and designed the experiments. M.B. performed all experiments. K.U. performed the computational analysis. S.C contributed to whole genome sequencing. K.U. Y.F., and M.B. wrote the manuscript. All authors reviewed the manuscript. All authors have read and agreed to the published version of the manuscript.

