## Supplementary Figures for "Restoration of the Functional *nif* Gene Cluster by Seven Excision and Two Inversion Events during Heterocyst Development in the Nitrogen-Fixing Cyanobacterium *Calothrix* sp. NIES-4101"

**Supplementary Figures: 12**

**Supplementary Figure S1.**	Growth of NIES-4101 in slant vials

**Supplementary Figure S2.**	*In silico* workflow for assembling reconstituted *nif* genes

**Supplementary Figure S3.**	PCR showing the presence of the restored *nif* genes

**Supplementary Figure S4.**	Multiple sequence alignments of NifH

**Supplementary Figure S5.**	Multiple sequence alignments of NifD

**Supplementary Figure S6.**	Multiple sequence alignments of NifK

**Supplementary Figure S7.**	Multiple sequence alignments of NifB

**Supplementary Figure S8.**	Conservation of gene arrangement of the *nif* gene cluster

**Supplementary Figure S9.**	Conservation of the *cis* element recognized by CnfR upstream of *nifB*

**Supplementary Figure S10.**	Alternative model for the reconstitution of the *nif* gene cluster

**Supplementary Figure S11.**	Models for the restoration of the four *nif*-related genes

**Supplementary Figure S12.**	Phylogenetic trees of 21 putative recombinases from NIES-4101

**Supplementary Tables: 6** (individual excel files)

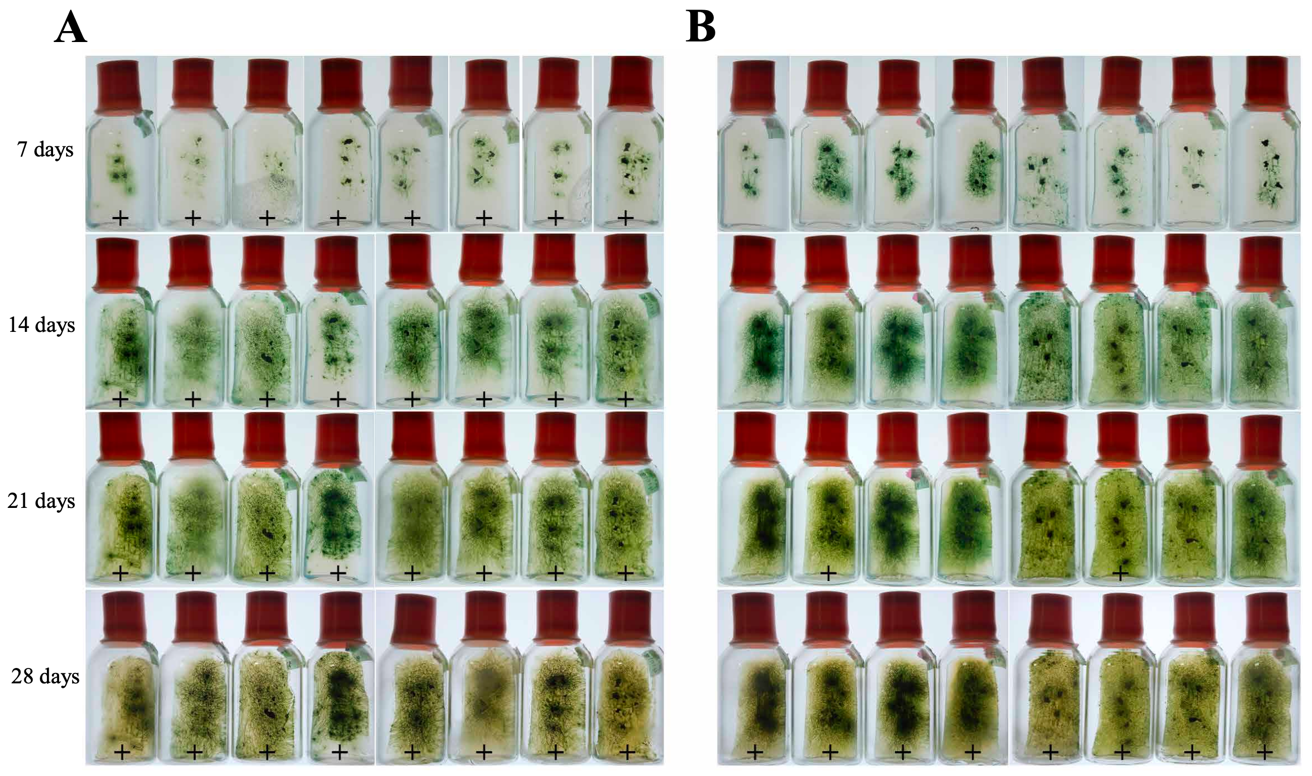

**Supplementary Figure S1.** Growth of NIES-4101 in slant vials for measurement of acetylene reduction activity. Due to the difficulty in suspending cells uniformly, clumps of NIES-4101 cells grown on a BG-11 agar plate were inoculated onto BG-11_0_ (**A**) or BG-11 (**B**) agar slant media and incubated under aerobic conditions at 30ºC (20 µmol m^-2^ s^-1^, continuous light). The acetylene reduction activity in each vial was measured after 7, 14, 21, and 28 days of incubation. Plus “+” marks indicate vials that showed acetylene reduction activity. The experiments were performed in eight replicates for both BG-11_0_ and BG-11 media.

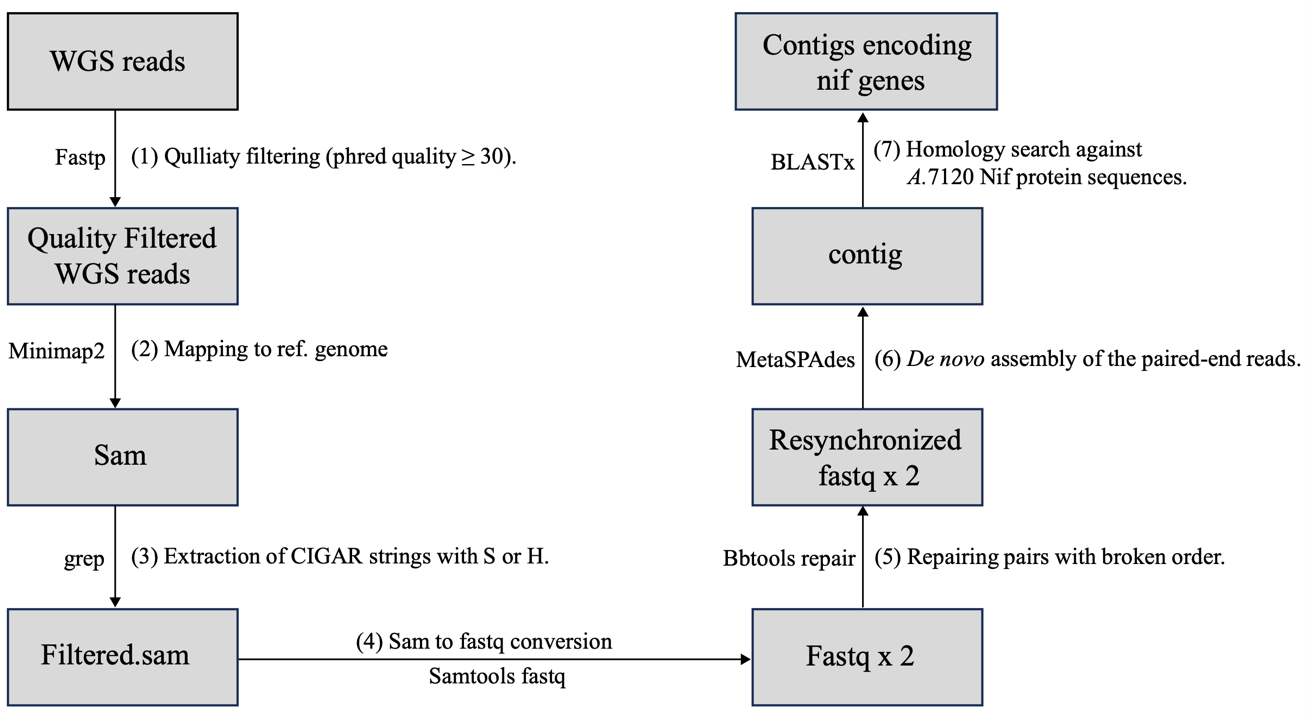

**Supplementary Figure S2.** *In silico* workflow for assembling reconstituted *nif* genes from bulk short read sequencing. The 177× paired-end whole-genome sequencing (WGS) reads were obtained on a G400 sequencer from DNA extracted from cells grown under nitrogen-fixing conditions (1). WGS reads were mapped to the NIES-4101 reference assembly (2); reads containing clipping in the alignment were extracted from the CIGAR in SAM (3,4,5) and assembled with a metagenome assembler (6). Among the resulting contigs, those corresponding to fragmented *nif* genes were extracted (7).

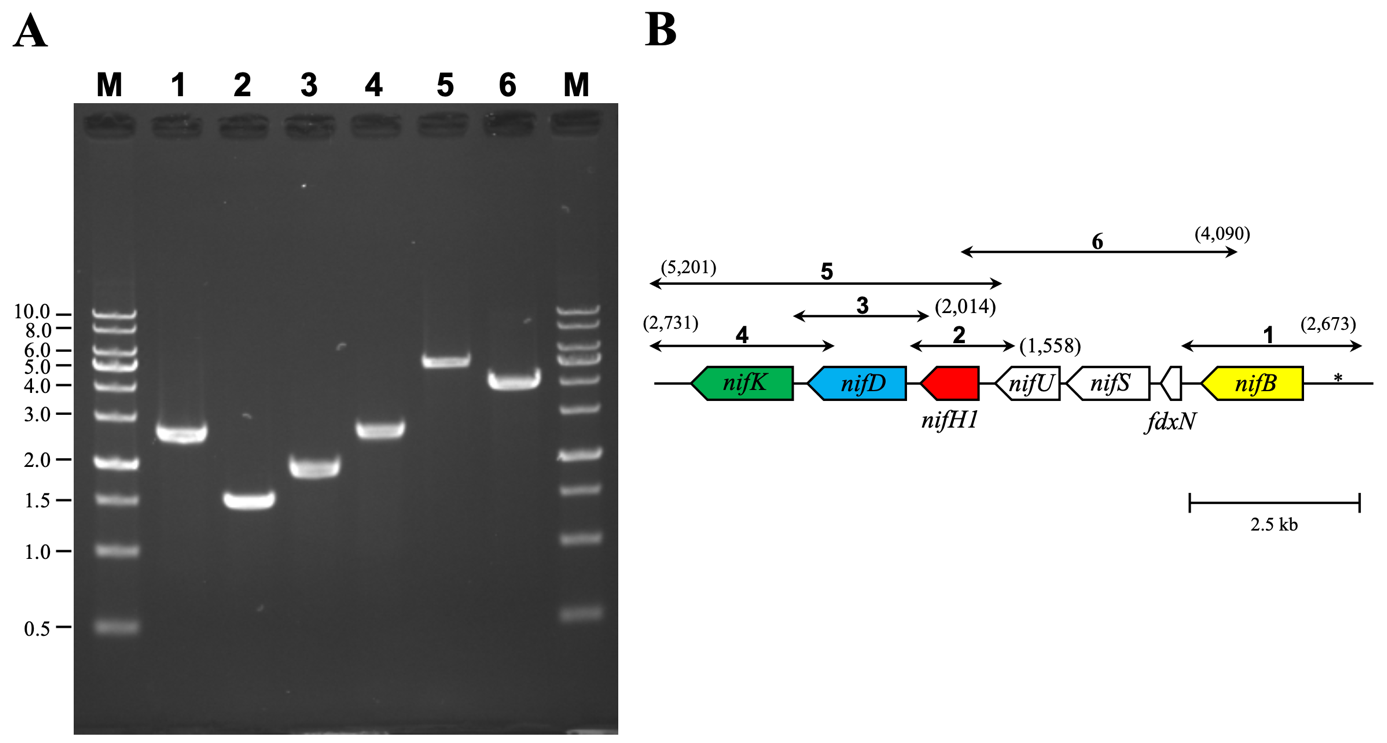

**Supplementary Figure S3.** PCR showing the presence of the restored *nif* genes. (**A**) PCR experiments were performed with six sets of primers designed for each restored gene (lanes 1 to 6). Genomic DNA extracted from cells grown on a BG-11_0_ agar plate for 2 weeks was used for the template. The 1-kb DNA ladder (same as Fig. 1B) was loaded on the lane M. (**B**) The fragments amplified with each primer set are shown by bars with arrows at both ends on the restored *nif* gene cluster. The number represents the lane number of panel **A**. The expected fragment size is shown in parentheses. The nucleotide sequences of the PCR primers used for this experiment are shown in Supplementary Tables S5 and S6. The asterisk represents the *cis* element in the *nifB* promoter.

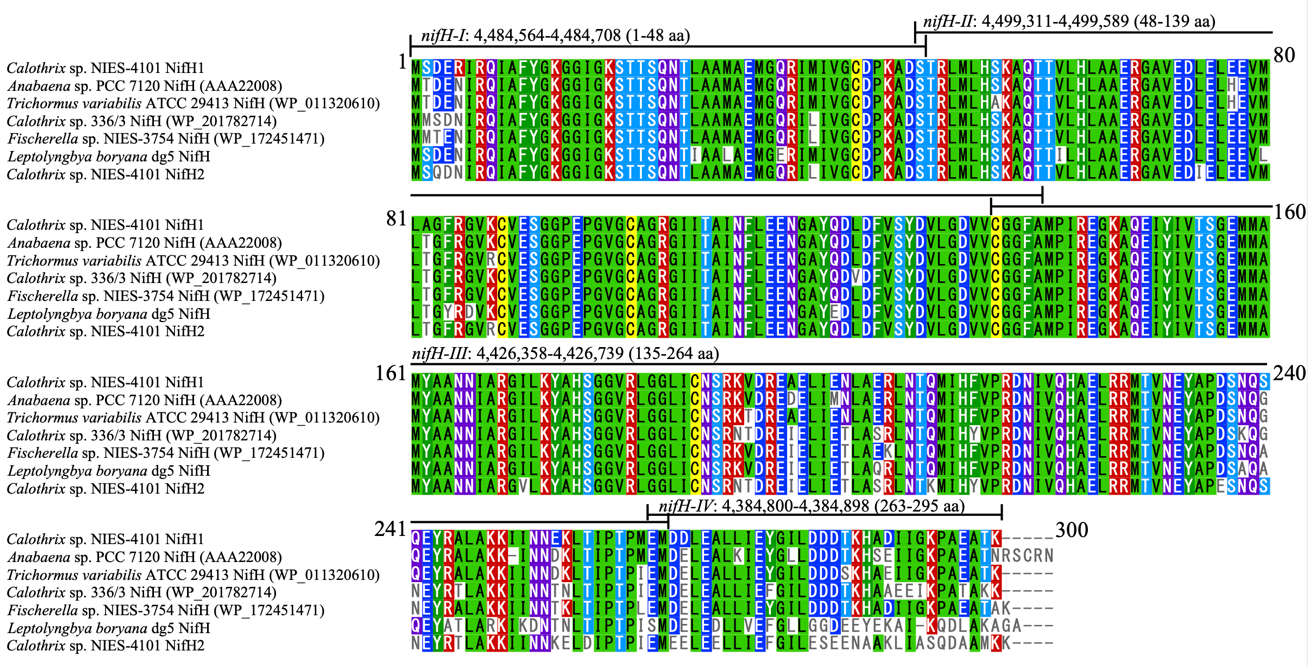

**Supplementary Figure S4.** Multiple sequence alignments of NifH. From top to bottom: 1) Restored NifH1 in NIES-4101, 2) *A.* 7120 NifH (accession: AAA22008), 3) *T. variabilis* ATCC 29413 NifH (accession: WP_011320610), 4) *Calothrix* sp. 336/3 NifH (accession: WP_201782714), 5) *Fischerella* sp. NIES-3754 NifH (accession: WP_172451471), 6) *L. boryana* dg5 NifH, and 7) NifH2 (from the monocistronic *nifH2* gene) in NIES-4101, respectively. Multiple sequence alignment (MSA) was constructed using MAFFT with the auto setting. The region encoded by each fragment (*nifH1*-I–IV) is indicated by horizontal bars.

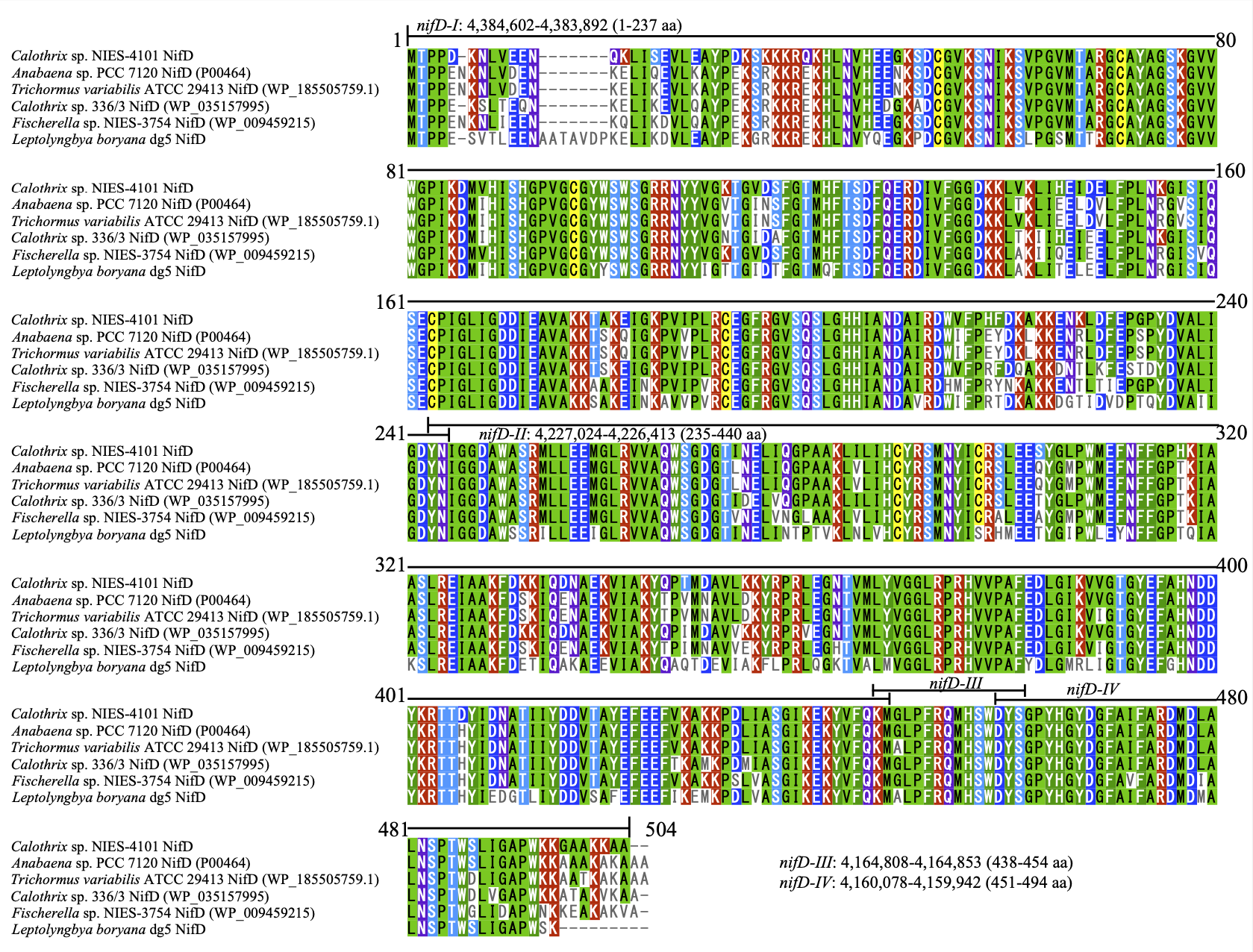

**Supplementary Figure S5.** Multiple sequence alignments of NifD. From top to bottom: 1) Restored NifD in NIES-4101, 2) restored NifD in strain *A.* 7120 (accession: P00464), 3) restored NifD in *T. variabilis* ATCC 29413 (accession: WP_185505759.1), 4) *Calothrix* sp. 336/3 NifD (009459215), 5) *Fischerella* sp. NIES-3754 NifD (accession: WP_009459215), and 6) *L. boryana* dg5 NifD, respectively. MSA was conducted using MAFFT with the auto setting. The region encoded by each fragment (*nifD*-I–IV) is indicated by horizontal bars.

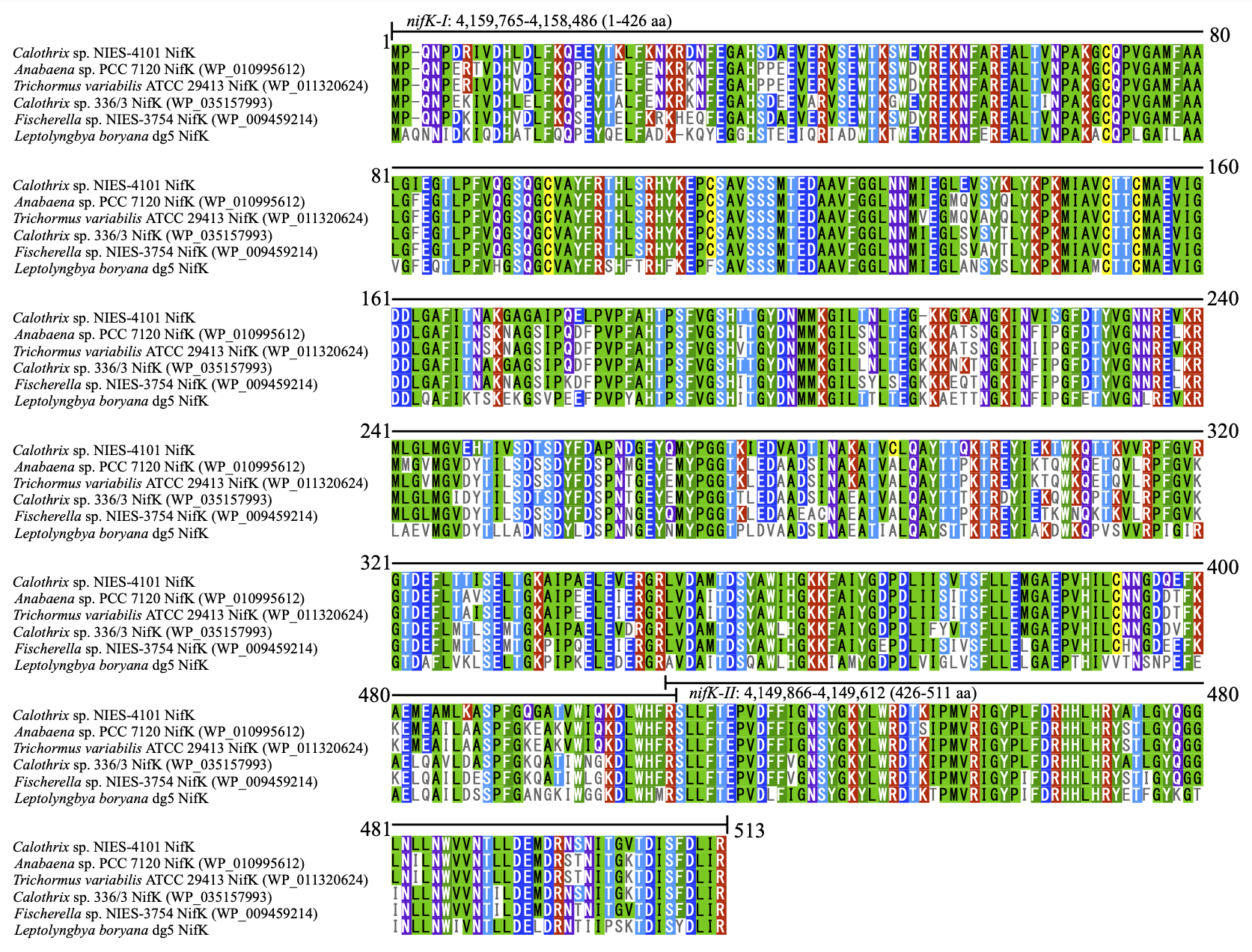

**Supplementary Figure S6.** Multiple sequence alignments of NifK. From top to bottom: 1) Restored NifK in NIES-4101, 2) *A.* 7120 NifK (accession: WP_010995612), 3) *T. variabilis* ATCC 29413 NifK (accession 011320624), 4) *Calothrix* sp. 336/3 NifK (accession: WP_035157993), 5) *Fischerella* sp. NIES-3754 NifK (accession: WP_009459214), 6) *L. boryana* dg5 NifK, respectively. MSA was conducted using MAFFT with the auto setting. The region encoded by each fragment (*nifK*-I–II) is indicated by horizontal bars.

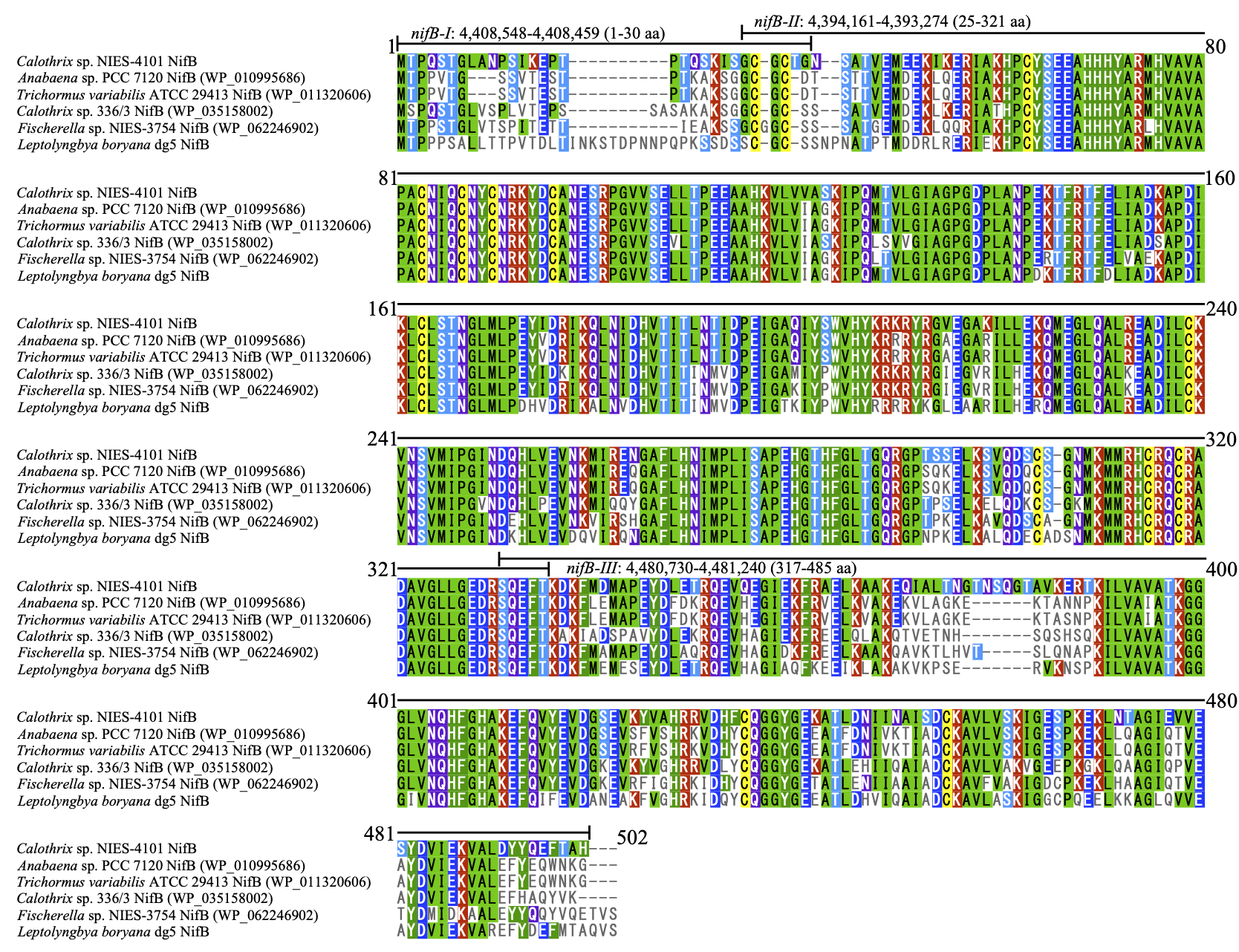

**Supplementary Figure S7.** Multiple sequence alignments of NifB. 1) Restored NifB in NIES-4101, 2) *A.* 7120 NifB (accession: WP_010995686), 3) *T. variabilis* ATCC 29413 NifB (accession: WP_011320606), 4) *Calothrix* sp. 336/3 NifB (accession: WP_35158002), 5) *Fischerella* sp. NIES-3754 NifB (accession: WP_062246902), 6) *L. boryana* dg5 NifB, respectively. MSA was conducted using MAFFT with the auto setting. The region encoded by each fragment (*nifB*-I–III) is indicated by horizontal bars.

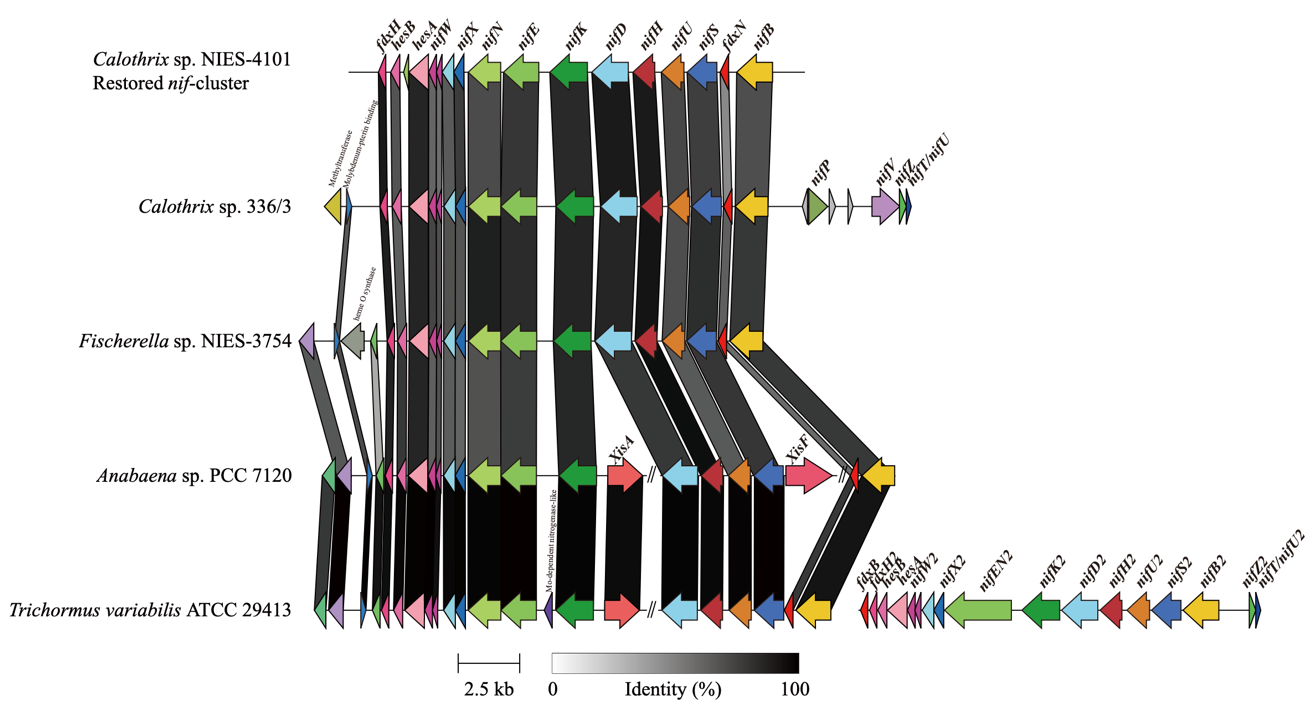

**Supplementary Figure S8.** Conservation of gene arrangement of the *nif* gene cluster between the restored *nif* cluster of NIES-4101 and four Nostocaceae species. From top to bottom: the restored *nif* cluster of NIES-4101, *Calothrix* sp. 336/3 and *Fischerella* sp. NIES-3754 (both of which have the complete *nif* gene cluster without any insertion sequences), *A*. 7120, and *T. variabilis* ATCC 29413. The *nifD* and *fdxN* genes in *A.* 7120 and the *nifD* gene in *T. variabilis* ATCC 29413 have the insertion sequences, which are indicated by “//”, and the *xisA* and *xisF* genes encoding recombinases in the insertion sequences are shown. The synteny plot was created using Clinker with the default settings (Gilchrist and Chooi 2021). Annotations for hypothetical proteins (two genes between *nifX* and *nifW*, one gene just downstream of *hesA*, one gene just downstream of *fdxH*, and two genes at the leftmost of *A.* 7120 and *T. variabilis* ATCC 29413) were omitted. Note that there is an extra *nif* gene cluster (*nif2*) in the *T. variabilis* ATCC 29413 genome shown in the right.

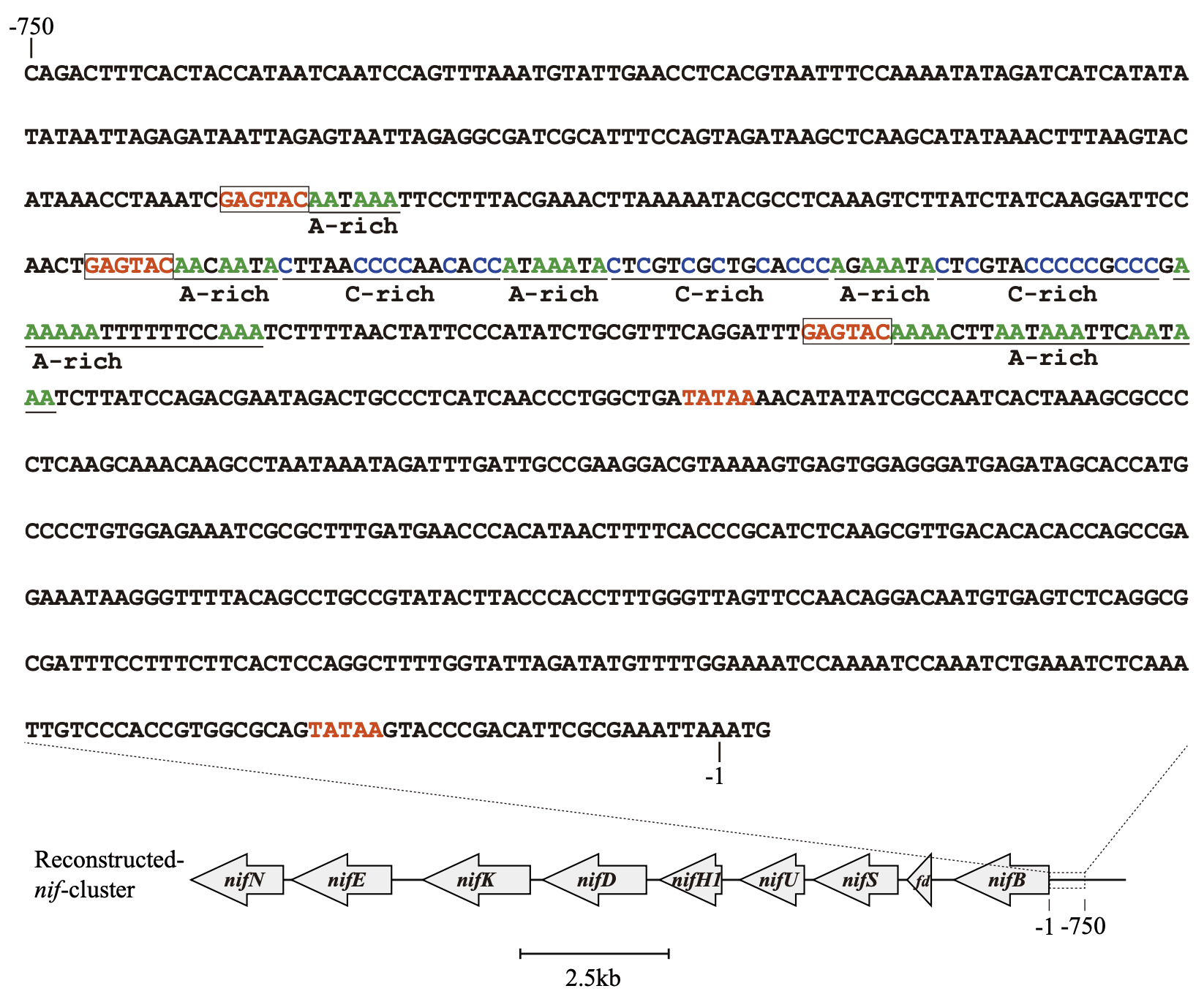

**Supplementary Figure S9.** Conservation of the *cis* element recognized by CnfR upstream of *nifB* in NIES-4101. Sequence elements for the recognition by CnfR (Tsujimoto *et al.* 2016) were found –1 to –750 upstream of the coding region of *nifB*. The C-rich regions, the A-rich regions, and the highly conserved GAGTAC motifs are shown in blue, green and red, respectively. The GAGTAC motifs are surrounded by boxes.

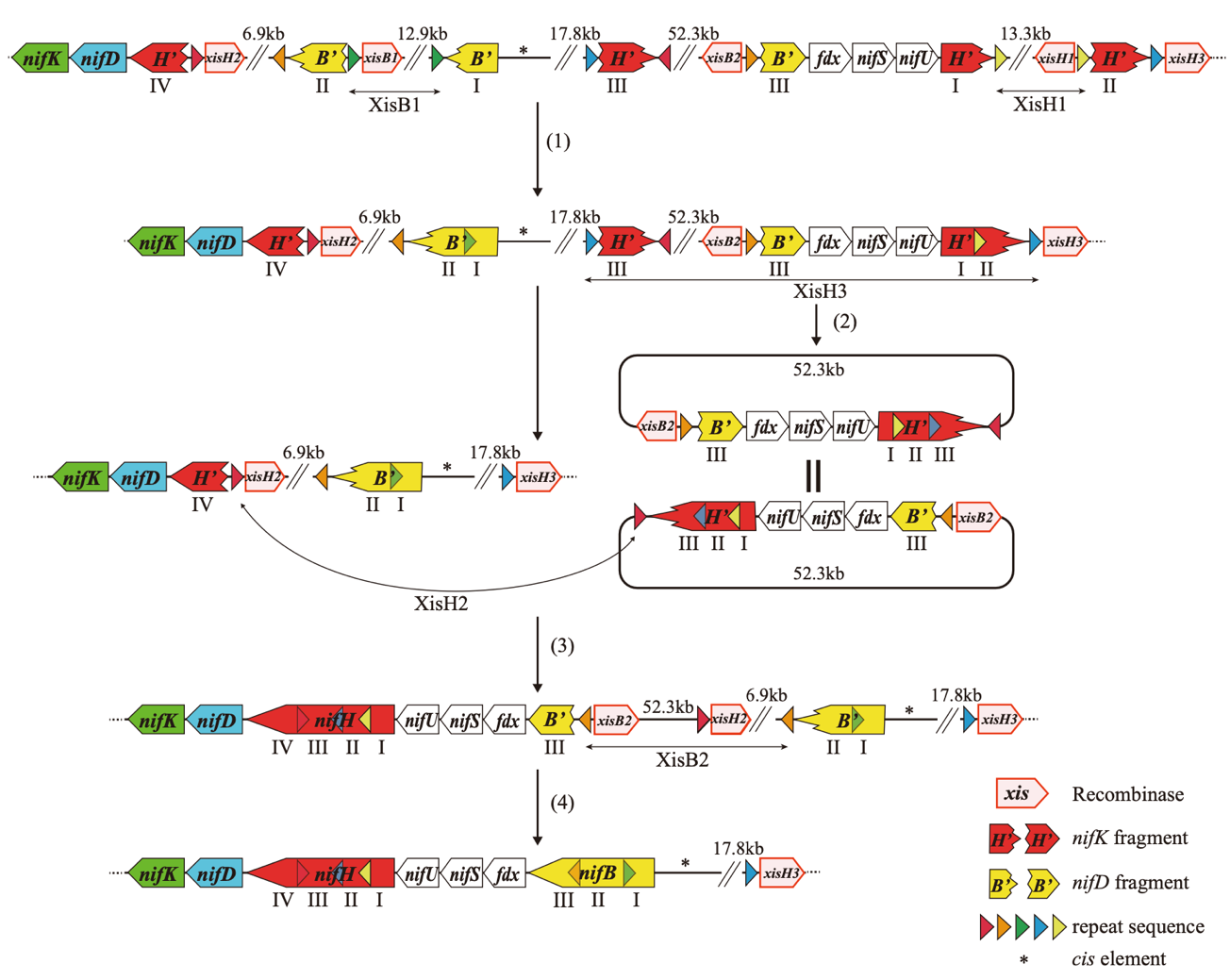

**Supplementary Figure S10.** Alternative model for the reconstitution of the *nif* gene cluster. (1) Sequences between direct repeats (black bars with arrowheads at both ends) are excised by site-specific recombinases (XisB1 and XisH1) same as Figure 4B. (2) The region from *nifH1*-III to *nifH1*-II is excised at the direct repeat site (blue triangles) by XisH3, resulting in the 58.6-kb circular double-stranded DNA (a transient episome). (3) The transient episome is reintegrated into the direct repeat site on the chromosome (red triangles) by XisH2. (4) The insertion sequence between the *nifB* fragments is excised by XisB2, forming the intact *nifB* gene same as Figure 4B. Irrelevant regions are indicated by “//”.

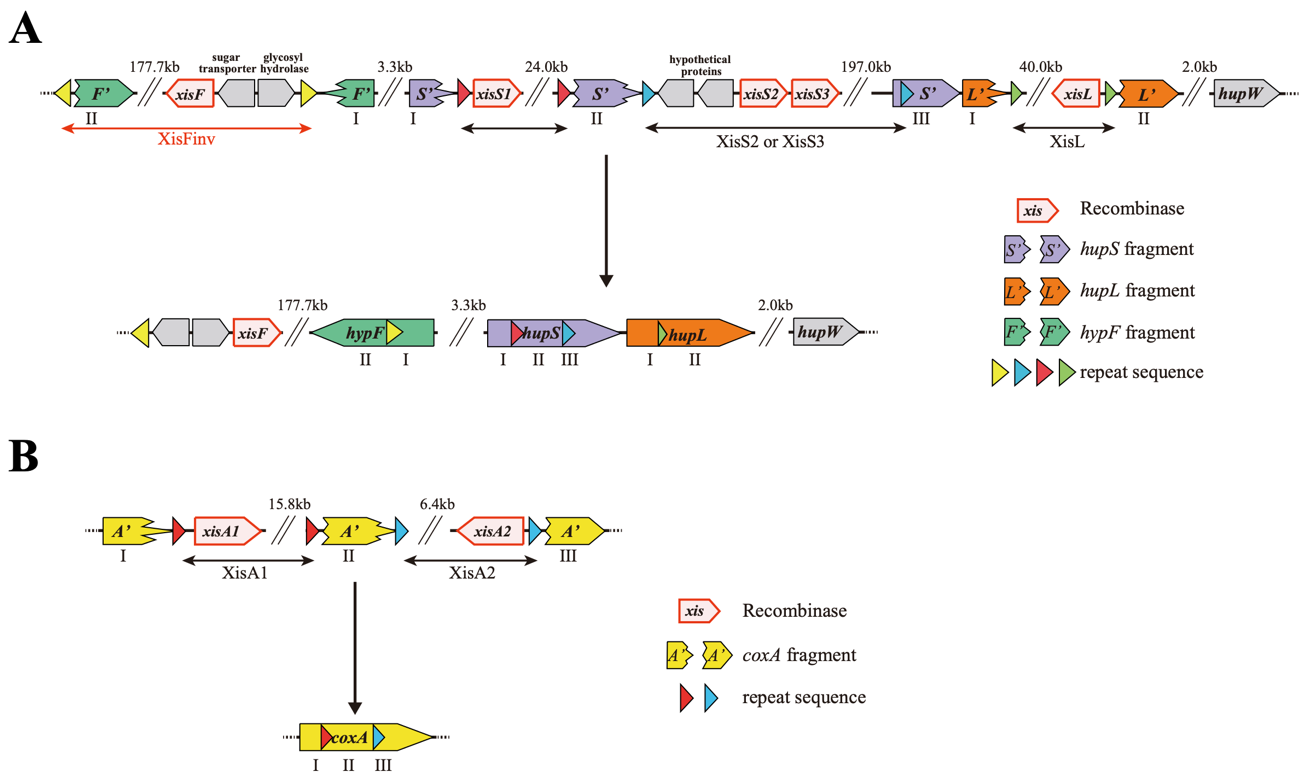

**Supplementary Figure S11.** Models for the restoration of the four *nif*-related genes. (**A**) A model for the restoration of the hydrogenase genes, *hypF* and *hupSL*. The full-length *hupS* and *hupL* genes are restored by three simple excision events (black bars with arrowheads at both ends). Two *hypF* fragments coded in the opposite directions are restored by the inversion event occurring between inverted repeats (yellow triangles, red arrowheads at both ends). The names of the recombinases involved in the excision and inversion are designated CnXisL, CnXisS1–3, and CnXisF after the gene names *hupL*, *hupS*, and *hypF*, respectively. Note that it is not known which of CnXisS2 and CnXisS3 are responsible for the excision event between *hupS*-II and *hupS*-III. (**B**) A model for the restoration of the *coxA* gene encoding subunit I of cytochrome *c* oxidase. The full *coxA* gene is reconstituted by two simple excision events (black bars with arrowheads at both ends). The names of the recombinases involved in the excision are designated CnXisA1 and CnXisA2.

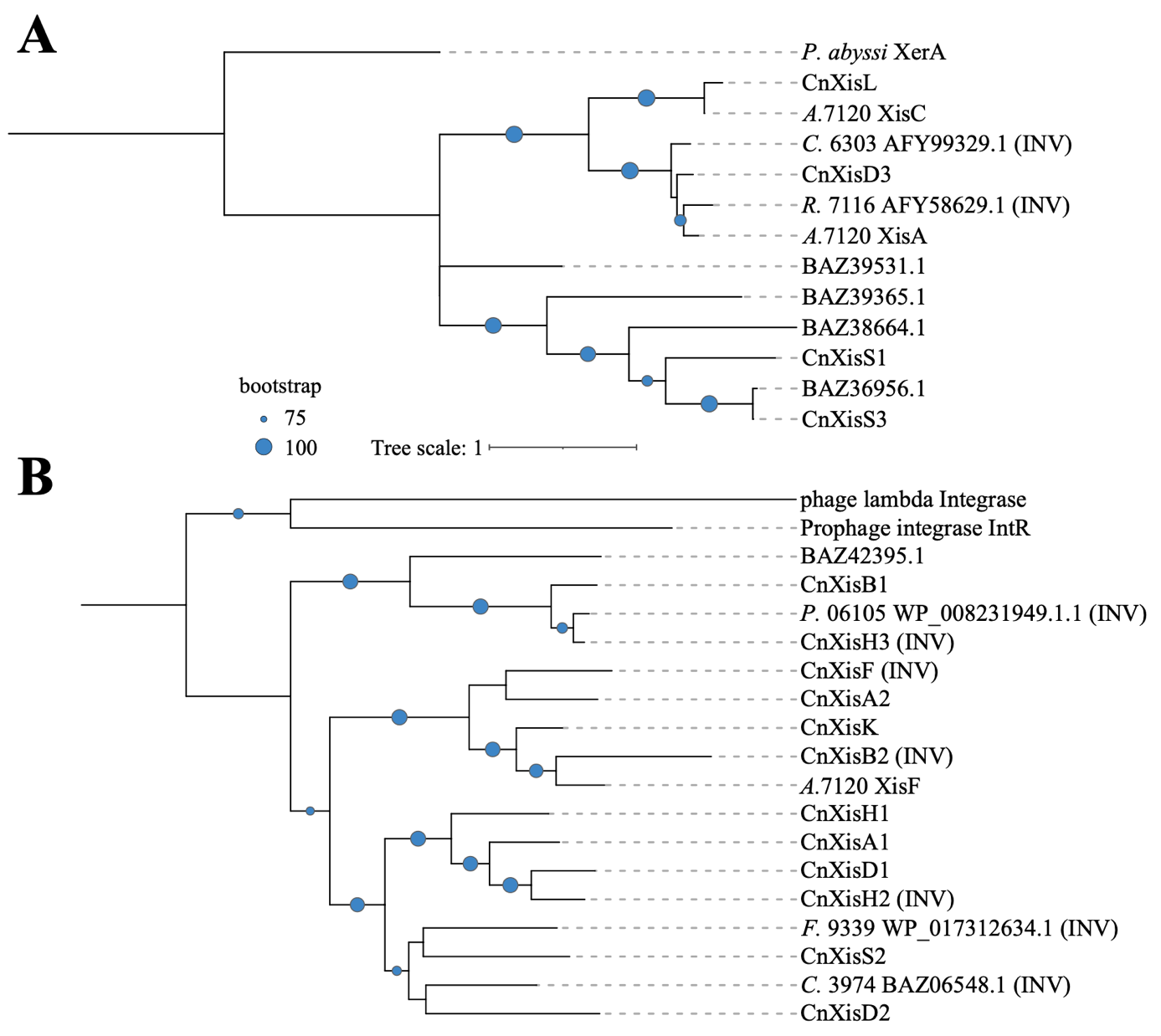

**Supplementary Figure S12.** Phylogenetic trees of 21 putative recombinases from NIES-4101. (**A**) Tyrosine recombinase family including 8 recombinases from NIES-4101, XisA and XisC from *A.* 7120, two putative recombinases from *Calothrix* sp. PCC 6303 (AFY99329.1) and *Rivularia* sp. PCC 7116 (AFY58629.1), and XerA from *Pyrococcus abyssi*, which shows structural similarity to XisA. (**B**) Serine recombinase family including 13 recombinases from NIES-4101, XisF from *A.* 7120, *E. coli* prophage integrase IntR (WP_000533640.1), three putative recombinases from *Plectonema* cf. *radiosum* LEGE 06105 (WP_008231949.1.1) and *Fischerella* sp. PCC 9339 (WP_017312634.1) and *Calothrix* sp. NIES-3974 (BAZ06548.1), and lambda phage integrase from Enterobacteria (WP_000040852.1), which shows structural identity with *A.* 7120 XisF. The recombinase catalyzing the putative inversion reaction is labelled (INV). MSAs were generated using MAFFT v7.08, and poorly conserved regions were trimmed using Clipkit v1.4.1 (Steenwyk et al. 2020) with “-m kpi-gappy” option. Phylogenetic inference was performed using the best-fit model in IQ-TREE v2.2.0.3 (LG + I + G4 for both A and B) (Minh et al. 2020) with ultra-fast 1,000 bootstrap approximation. The interactive web-based tree visualization service iTOL v6 (Letunic and Bork, 2021) was used to render the phylogenetic tree.
